# Structural basis for the hyperthermostability of an archaeal glutaminase induced by post-translational succinimide formation

**DOI:** 10.1101/2021.03.02.433506

**Authors:** Aparna Vilas Dongre, Sudip Das, Asutosh Bellur, Sanjeev Kumar, Anusha Chandrashekarmath, Tarak Karmakar, Padmanabhan Balaram, Sundaram Balasubramanian, Hemalatha Balaram

## Abstract

Stability of proteins from hyperthermophiles enabled by reduction of conformational flexibility is realized through various mechanisms. Presence of a stable, hydrolysis-resistant succinimide arising from cyclization of the side chains of aspartyl/asparaginyl residues with backbone amide -NH of the succeeding residue would restrain the torsion angle Ψ. Here, we describe the crystal structure of *Methanocaldococcus jannaschii* glutamine amidotransferase (MjGATase) and address the mechanism of a succinimide-induced increased thermostability using molecular dynamics simulations. This study reveals the interplay of negatively charged electrostatic shield and n→π* interactions in preventing succinimide hydrolysis. The stable succinimidyl residue induces formation of a ‘conformational-lock’, reducing protein flexibility. Protein destabilization upon replacement with the Φ-restricted prolyl residue highlights the specificity of the conformationally restrained succinimidyl residue in imparting hyperthermostability. The conservation of succinimide-forming tripeptide sequence (E(N/D)(E/D)) in a group of archaeal GATases suggests an adaptation of this otherwise detrimental post-translational modification as an inducer of thermostability.

## Introduction

Succinimide formation by intramolecular cyclization at Asx/Xxx (Asp/Asn, Gly) segments in proteins is an intermediate step in the process of deamidation, which often accompanies aging(*1*). Succinimidyl/aminosuccinyl (Asu, succinimide or SNN) residues are unstable, readily hydrolysing under aqueous conditions to Asp and β-Asp (isoAsp) residues(*2*–*4*). Facile stereo inversion at the C^α^ atom of the succinimide results in the formation of D-Asp, often used as a marker of aging(*4*). Stable succinimides are uncommon in proteins with relatively few well characterised examples(*5*–*9*). We recently established, mass spectrometrically, the presence of a stable succinimide in the hyperthermophilic protein glutamine amidotransferase, a subunit of the enzyme GMP synthetase (GMPS) from *Methanocaldococcus jannaschii*(*10*). This post-translational modification at Asn109 confers remarkable thermal stability on the WT protein, which resists unfolding up to 100 °C. Mutants lacking the succinimide melted at appreciably lower temperatures(*10*). Many structural features have been implicated in conferring stability to proteins from thermophilic organisms. These include, shortening or deletion of loops(*11*), increased occurrence of proline residues(*12, 13*), increased number of hydrogen bonds and disulphide bridges(*14*–*16*) and post-translational modifications like glycosylation(*17*) and methylation(*18*). The observation of succinimide induced hyper thermostability in *M. jannaschii* glutamine amidotransferase (MjGATase) prompted us to investigate the structural origins of this phenomenon. Towards this end, we determined the crystal structure of MjGATase in both *apo* and ligand bound forms, revealing that the succinimide residue (SNN109) lies at the solvent exposed tip of a loop in the protein, with the succeeding residue Asp(D)110 shielding its one face from hydrolytic attack by water, while its other face is shielded by the backbone carbonyl group of the preceding residue Glu(E)108 through n→π* interaction. Structure determination of the D110G mutant showed an absence of electron density for residues 108-110, suggesting enhanced flexibility in the absence of the succinimide constraint. In contrast, the structure of N109P mutant showed that the absence of key SNN mediated interactions present in the WT protein could be the cause for impaired thermostability of the mutant. To probe the origins of protein stabilization by the succinimidyl residue, we turned to molecular dynamics simulation (MDS), which has been widely used to delineate functional dynamics of proteins(*19*–*23*), mechanisms of protein folding/unfolding pathways(*24*–*30*) and protein stability mechanisms(*27*–*33*). MDS coupled with replica exchange with solute(enzyme) scaling (REST2)(*34*), a protocol that mitigates the problem of insufficient sampling of conformational space, and well-tempered metadynamics (WT-MTD)(*35*) simulations have been used to study the difference in the thermal stabilities of the WT enzyme and the single site mutants N109S and N109P (referred to as SNN109S and SNN109P hereafter), which lack the succinimide. REST2 simulations highlight the role of a series of backbone hydrogen bonds facilitated by the succinimide constraint, which imposes a local conformational-lock. Additional long-range side-chain hydrogen bonds and van der Waals interactions involving residues 109 and 110, contribute to succinimide induced stabilisation. Additionally, well-tempered meta-dynamics (WT-MTD) simulations on WT and SNN109S mutant reinforce our observation on succinimide-induced conformational-lock.

## Results and Discussion

### Analysis of the structure of MjGATase and site-specific mutants

Crystals of MjGATase (Space group P32, 2 molecules/asymmetric unit) yielded structures of both apo and ligand (acivicin) bound forms, to a resolution of 1.67Å, by molecular replacement (MR), using *Pyrococcus horikoshii* GATase (PhGATase, PDB ID 1WL8)^34^ as a model. Data collection and refinement statistics are reported in Table 1. The overall fold of the molecule and the position of the segment with respect to the succinimidyl residue along with the electron density corresponding to this post-translational modification are shown in Figure 1a. A common feature of the class I amidotransferases is the central region comprising of seven β strands surrounded by α helices and loops(*36*). This structural feature is conserved across eukaryotes, prokaryotes and archaea, as seen from the superposition of the structure of MjGATase with GATases of GMPS from seven representative organisms (Supplementary Figure 1a). The succinimide containing segment, residues 108-110 (Glu-Asu(SNN)-Asp), lies at the solvent exposed tip of the loop (residues 106-114) connecting the two antiparallel strands (residues 97-105 and 115-124) of a β-hairpin (Fig. 1c). The segment 108-110 forms a local α-turn structure stabilized by a 13-atom hydrogen bond, Glu108 CO ---HN Phe112 (Fig. 1b)(*37*–*39*). The catalytic residues, Cys76, His163 and Glu165 are proximate to the succinimide containing loop. Active site occupancy is without any effect on the succinimide containing loop as established by the determination of the structure of an acivicin (5CS) modified enzyme (Supplementary Figure 1b). Acivicin is a glutamine analogue which irreversibly modifies the active site cysteine residue(*40*).

**Table 1:**
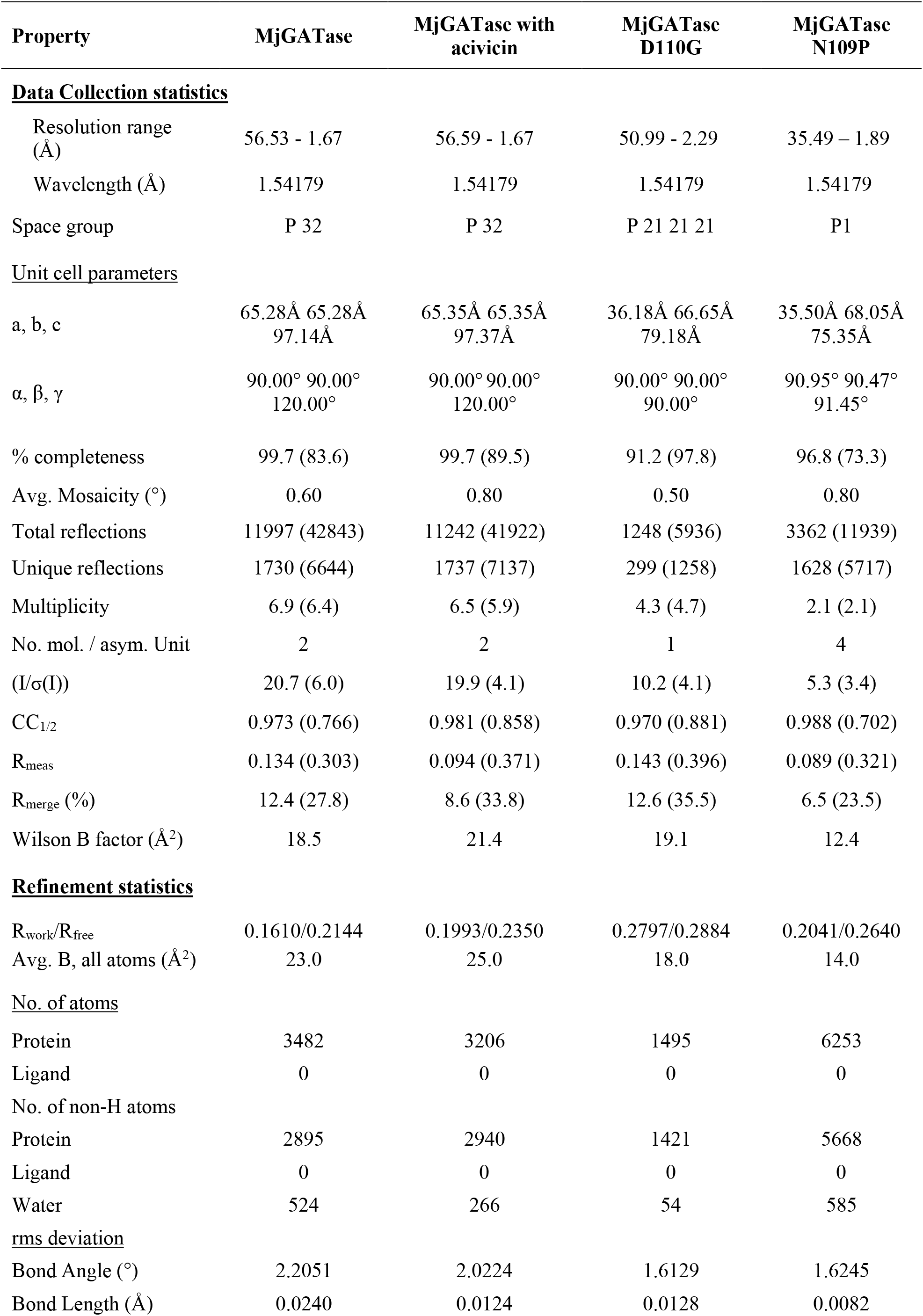
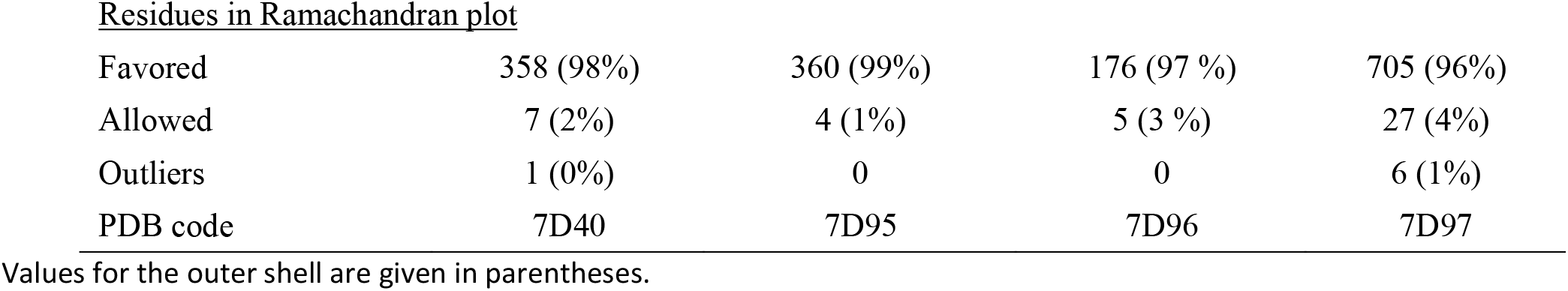
Summary of X-ray crystallographic data-collection and refinement statistics.

| Property | MjGATase | MjGATase with<br>acivicin | MjGATase<br>D110G | MjGATase<br>N109P |
| --- | --- | --- | --- | --- |
| **<u>Data Collection statistics</u>** |  |  |  |  |
| Resolution range<br>(Å) | 56.53 - 1.67 | 56.59 - 1.67 | 50.99 - 2.29 | 35.49 – 1.89 |
| Wavelength (Å) | 1.54179 | 1.54179 | 1.54179 | 1.54179 |
| Space group | P 32 | P 32 | P 21 21 21 | P1 |
| **<u>Unit cell parameters</u>** |  |  |  |  |
| a, b, c | 65.28Å 65.28Å<br>97.14Å | 65.35Å 65.35Å<br>97.37Å | 36.18Å 66.65Å<br>79.18Å | 35.50Å 68.05Å<br>75.35Å |
| $\alpha, \beta, \gamma$ | 90.00° 90.00°<br>120.00° | 90.00° 90.00°<br>120.00° | 90.00° 90.00°<br>90.00° | 90.95° 90.47°<br>91.45° |
| % completeness | 99.7 (83.6) | 99.7 (89.5) | 91.2 (97.8) | 96.8 (73.3) |
| Avg. Mosaicity (°) | 0.60 | 0.80 | 0.50 | 0.80 |
| Total reflections | 11997 (42843) | 11242 (41922) | 1248 (5936) | 3362 (11939) |
| Unique reflections | 1730 (6644) | 1737 (7137) | 299 (1258) | 1628 (5717) |
| Multiplicity | 6.9 (6.4) | 6.5 (5.9) | 4.3 (4.7) | 2.1 (2.1) |
| No. mol. / asym. Unit | 2 | 2 | 1 | 4 |
| (I/$\sigma$(I)) | 20.7 (6.0) | 19.9 (4.1) | 10.2 (4.1) | 5.3 (3.4) |
| CC<sub>1/2</sub> | 0.973 (0.766) | 0.981 (0.858) | 0.970 (0.881) | 0.988 (0.702) |
| R<sub>meas</sub> | 0.134 (0.303) | 0.094 (0.371) | 0.143 (0.396) | 0.089 (0.321) |
| R<sub>merge</sub> (%) | 12.4 (27.8) | 8.6 (33.8) | 12.6 (35.5) | 6.5 (23.5) |
| Wilson B factor (Å<sup>2</sup>) | 18.5 | 21.4 | 19.1 | 12.4 |
| **<u>Refinement statistics</u>** |  |  |  |  |
| R<sub>work</sub>/R<sub>free</sub> | 0.1610/0.2144 | 0.1993/0.2350 | 0.2797/0.2884 | 0.2041/0.2640 |
| Avg. B, all atoms (Å<sup>2</sup>) | 23.0 | 25.0 | 18.0 | 14.0 |
| **<u>No. of atoms</u>** |  |  |  |  |
| Protein | 3482 | 3206 | 1495 | 6253 |
| Ligand | 0 | 0 | 0 | 0 |
| **<u>No. of non-H atoms</u>** |  |  |  |  |
| Protein | 2895 | 2940 | 1421 | 5668 |
| Ligand | 0 | 0 | 0 | 0 |
| Water | 524 | 266 | 54 | 585 |
| **<u>rms deviation</u>** |  |  |  |  |
| Bond Angle (°) | 2.2051 | 2.0224 | 1.6129 | 1.6245 |
| Bond Length (Å) | 0.0240 | 0.0124 | 0.0128 | 0.0082 |
| Residues in Ramachandran plot |  |  |  |  |
| Favored | 358 (98%) | 360 (99%) | 176 (97 %) | 705 (96%) |
| Allowed | 7 (2%) | 4 (1%) | 5 (3 %) | 27 (4%) |
| Outliers | 1 (0%) | 0 | 0 | 6 (1%) |
| PDB code | 7D40 | 7D95 | 7D96 | 7D97 |

Values for the outer shell are given in parentheses.

**Figure 1:**
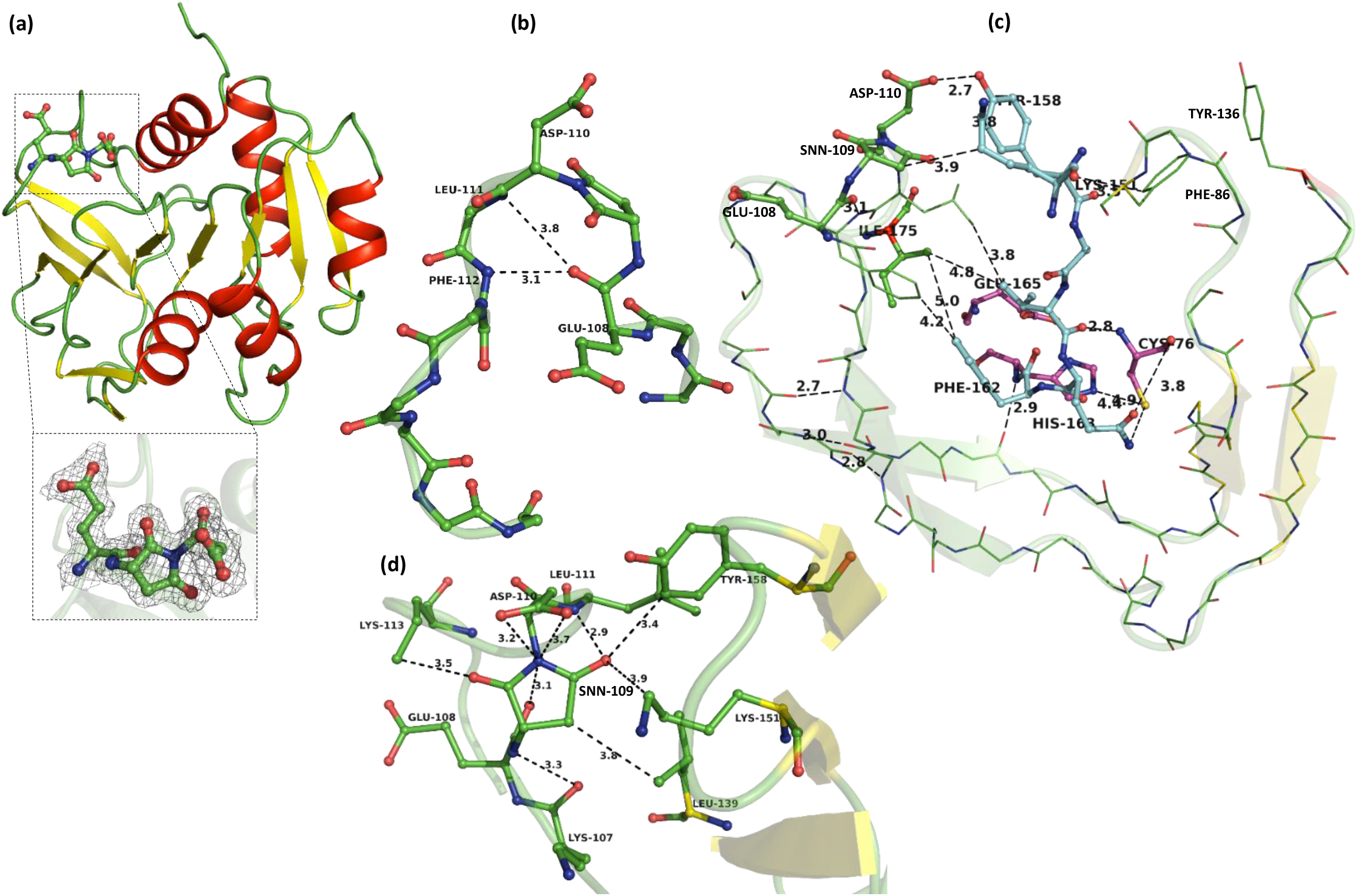
Interactions of SNN109 in MjGATase. (a) Structure of MjGATase. Inset: Succinimidyl residue with electron density (2F_0_-F_c_, contoured to 1σ). (b) SNN-harbouring loop bends to form overlapping β-turn (3.8 Å H-bond) and α-turn (3.1 Å H-bond). (c) SNN is connected to active site and distal β-sheet residues through tertiary contacts. SNN-containing β-hairpin residues, green; active site residues, purple and residues mediating tertiary contacts, cyan. (d) Residues in the vicinity of SNN (4 Å distance cut-off).

The network of contacts illustrated in Figure 1c highlight close interactions between residues flanking the succinimide moiety and the segment 151-165 containing two active site residues. Key inter-segment contacts are the hydrogen bonds between the side-chains of Asp110 and Tyr158 and the van der Waals contacts between SNN109 and the side-chains of Lys151, Leu111 and Val160, and Phe112 and Phe162 (Supplementary Table 1). The stabilizing effect of the succinimide on the temperature dependence of MjGATase activity has been previously noted with the WT enzyme showing stability and activity up to about 100 °C, while mutants lacking this PTM lose activity by about 85 °C(*10*). The unusual stability of the succinimide residue may be rationalised by the network of contacts shown in Figure 1d. The carboxymethyl side-chain of Asp110 folds over one face of the five membered succinimide ring, while the carbonyl group of Glu108 protects the opposite face (Fig. 1d and 3a). Interestingly, mutation of Asp110 to Gly increases the flexibility of this tripeptide segment resulting in the absence of electron density for the residues E108-G110 while rest of the structure overlaps well with that of WT (Supplementary Figure 1c, d). The Glu108 CO---SNN109 CO contacts of 2.7 and 3.3 Å lie within the ranges established by Raines and co-workers for stabilizing n→π* interactions in biomolecules(*41*–*43*). An overall stabilization energy of 11.7 kJ/mol is estimated from natural bond orbital (NBO) analysis(*44*), for the interaction of Glu108 CO with the CO groups of the succinimide (Fig. 3b, c and Supplementary Figure 2).

Removal of the succinimide induced conformational restraint in the SNN109S mutant results in reduction of stability. Introduction of a different restraint in the form of a prolyl residue at position 109 in the SNN109P mutant resulted in greater loss of stability. As expected, the WT and mutants showed similar far-UV CD spectra at 25 °C (Fig. 2a), but a variation in thermal stability was clearly evident; WT being stable at 100 °C while SNN109P exhibited a T_m_ of 78 °C, the lowest of all the mutants studied (Fig 2b). Notably, a search of the sequence database yielded 84 non-redundant archaeal sequences, belonging to either Thermococci or Methanococci, in which a very high degree of conservation was noted for residues 108-110, 139, 151 and 158 (numbering follows MjGATase) (Supplementary Figure 4 a, b). Although the overall structure of MjGATase_SNN109P mutant overlays well on the WT structure with an RMSD of 0.390 Å (Supplementary Figure 3 a), the SNN loop shows a movement of the backbone that is reflected as a large change in the Φ, Ψ values of residues 107, 108 and 109 (Fig. 2 c, d, and Supplementary Figure 3 b). As a consequence, H-bonds involving Glu108 CO are absent in MjGATase_SNN109P leading to the loss of α and β-turns (Fig. 2e) but the polypeptide chain in MjGATase_SNN109P bends in a manner leading to H-bonds between Pro109 CO---Phe112 NH (3.0 Å, β-turn), and Asp110 CO---Lys113 NH (3.1 Å, β -turn) resulting in the formation of two consecutive β-turns (Fig. 2f). Further, due to the different positioning of the side-chain of Glu108, the salt bridge with Arg117 is broken in the mutant (Supplementary Figure 3c).

**Figure 2:**
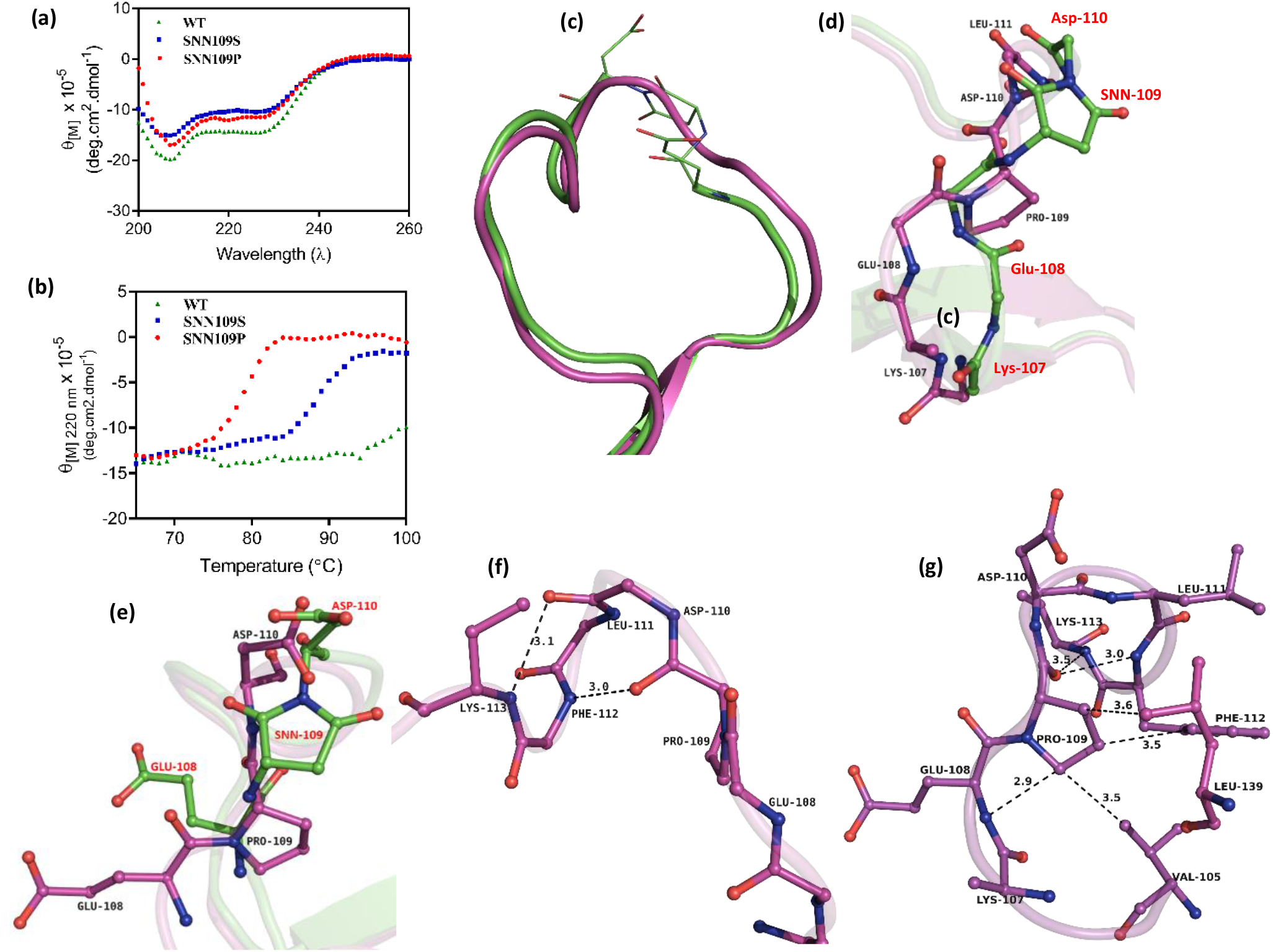
Impact of replacement of SNN with prolyl residue. (a) Structural differences between WT (green) and SNN109P (purple) in loop region (residues 105-116). (b) Different paths traced by polypeptide 107-111. (c) Larger difference observed in conformation of side-chains of residues 108-110. (d) α, β turns in WT (Figure 1, panel b) replaced with two consecutive β-turns formed by H-bonding between Pro109 CO---Phe112 NH and Asp110 CO---K113 NH in MjGATase_SNN109P. (e) MjGATase_SNN109P structure revealed loss of specific SNN-mediated contacts. (f) Far-UV CD spectra show secondary structure conservation at 25 °C. (g) Molar ellipticity illustrates loss of secondary structure at higher temperatures for mutants.

Pro109 shows overall fewer interatomic interactions compared to SNN109 (Fig. 2g) including loss of H-bond between K107 CO ----NH of N109 (SNN) present in the WT, owing to the imino nitrogen of the prolyl residue. Apart from these local changes, the movement of the backbone in MjGATase_SNN109P has also resulted in loss of tertiary interactions of the SNN loop with the protein core. In the mutant, the electron density for the side-chain of Lys107 is absent whereas the backbone geometry in the WT permits the formation of a salt bridge between Lys107 and Glu137. The interaction between Lys107 NH---Leu139 CO (3.1 Å in the WT) and van der Waals interactions of SNN with K151 and Y158 are lost in the mutant. In addition, the movement of Asp110 side-chain COOH has resulted in a weakening of the interaction with Tyr158 OH (Supplementary figure 3 d). Taken together, the absence of these interactions has dramatically reduced the stability of the mutant.

### Hydrolytic Stability of the Succinimide

Hydrolysis of the succinimide requires approach of a water molecule to either of its carbonyl group in a direction perpendicular to the plane of the ring(*45, 46*). Both faces of the ring are sterically shielded from solvent approach, with the Asp110 side-chain carboxylate lying parallel to one face while the other is shielded by the Glu108 backbone carbonyl group (Fig. 3a). However, at elevated temperatures, movement of the Asp110 side chain due to population of alternative rotamers may permit the close approach of a water molecule. Three water molecules are observed in the vicinity of the succinimide in the crystal structure (Fig. 3d). Notably, all three molecules are conserved in the structure of the *P. horikoshii* enzyme (PhGATase, PDB: 1WL8). None of the water molecules meet the orientational requirements for hydrolytic attack. We therefore turned to MD simulations as a function of temperature to examine the possibility of side-chain and water reorientation in the vicinity of the succinimide. Figure 3e defines the angular parameters θ1 and θ2 and the distances r1 and r2 which describe the Asp110 side-chain orientation and the approach of the nucleophile to either of the succinimide carbonyl groups. MD simulation shows retention of the parallel orientation of Asp110 side-chain carboxylate group over succinimide plane even at elevated temperature (Supplementary Figure 5b, Supplementary Movie1).

**Figure 3:**
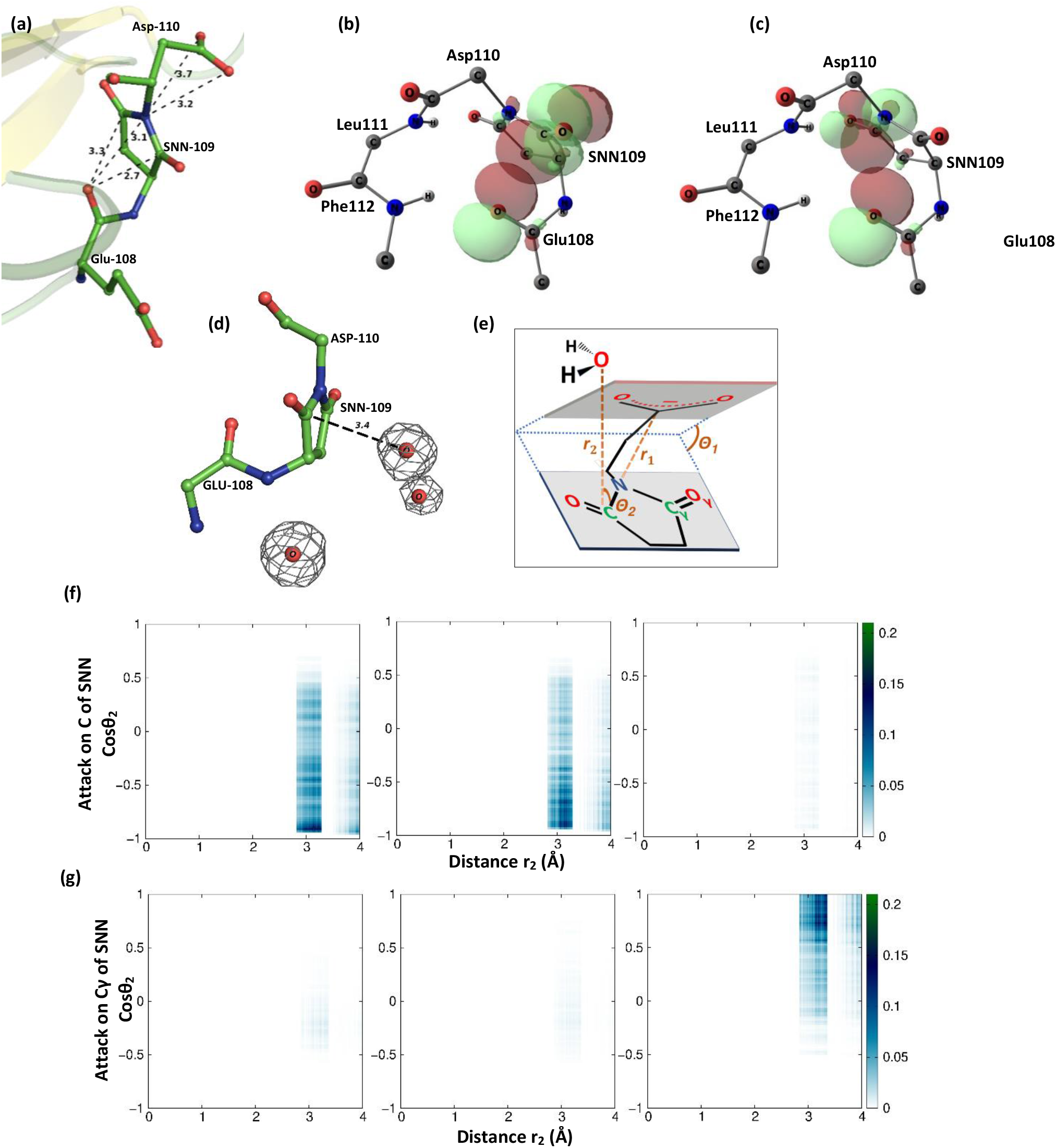
Shielding of SNN from hydrolysis. (a) Side-chain of Asp110 shields front-end of SNN from hydrolysis. SNN is shielded from rear-end through n→π* interactions between the backbone carbonyl oxygen of Glu108 to backbone (b) and side chain (c) carbonyl carbons of SNN 109. Except for SNN, only backbone atoms (NH-C_α_-CO) for rest of the residues are shown(*47*). (d) Three water molecules in the vicinity of SNN in the crystal structure of MjGATase are present in the structure of PhGATase. (e) Schematic representation of the distances and angles used in panels (f) and (g). θ_1_ is angle between side-chain carboxylate plane of Asp 110 (C_γ_-O_δ1_-O_δ2_ plane) and succinimide plane (C-N-Cγ plane). r_1_ is distance between N and C_γ_ atoms of Asp 110 (i.e., distance between these two planes). r_2_ represents distance of oxygen of water molecules within 4 Å of C (or Cγ) of SNN and θ_2_ is the angle between SNN-plane and the line joining this C (or Cγ) atom to such oxygen atoms. Probability distributions P(r_2_, cosθ_2_) showing the approach of water towards C (f) and Cγ (g) atoms of SNN.

This rigid orientation of the side-chain carboxylate group of D110 arises from its strong hydrogen bond with the side-chain hydroxyl oxygen of Tyr158 from one of the core β-strands of MjGATase (Fig. 1c and Table 2). Thus, irrespective of temperature, water hardly approaches either of the succinimide carbonyl groups in a direction perpendicular to the SNN ring plane (Fig. 3f, 3g). In MD simulations of the homologous enzyme PhGATase, the parallel orientation of the side-chain carboxylate plane of the succeeding Glu (E113) residue shields the succinimide from the hydrolytic attack in a plane perpendicular to one face of the succinimide ring (Supplementary Figure 5c, 5d). However, in MjGATase_D110G, the absence of Asp side-chain following the succinimide residue allows the approach of water in a direction perpendicular to the succinimide plane (Supplementary Figure 5e), leading to the facile hydrolysis of the succinimide in this mutant. These findings reaffirm our experimental observation that Asp110 plays an important role in the stabilization of succinimide(*10*). Taken together, the experiments and MD simulations point to a major role for the Asp110 residue in stabilizing the succinimide modification at position 109.

**Table 2:** Average number of hydrogen bond(s) controlling the stability of succinimide and describing the conformational lock in the SNN-loop region from MD simulations. Average number of backbone hydrogen bonds between two β-strands forming the distal β-sheet connected to the β-hairpin containing succinimide are also reported. For WT-MTD, the position of free energy minimum with respect to radius of gyration (R_g_) of residues 108-115 are mentioned within parenthesis and the average number of hydrogen bonds are calculated over configurations around the corresponding free energy minimum (for WT: 5.36-5.56 Å; for SNN109S mutant, lower free energy minimum: 5.52-5.72 Å; for SNN109S mutant, higher free energy minimum: 6.90-7.10 Å). A donor-acceptor distance cut-off of 3.5 Å and donor-acceptor-hydrogen angle cut-off of 40° has been used to define a hydrogen bond. *Number of hydrogen bonds in the X-ray crystal structure of WT MjGATase was calculated based on only donor-acceptor distance criterion. **For crystal structure, it is not an average number.

| Type | Atom pair(s) | Average* number of hydrogen bond(s) |  |  |  |  |  |  |  |  |  |  |  |  |
| --- | --- | --- | --- | --- | --- | --- | --- | --- | --- | --- | --- | --- | --- | --- |
|  |  | ** | 27 °C |  |  | 70 °C |  |  | 87 °C |  |  | WT-MTD (27 °C) |  |  |
|  |  |  | WT | SNN 109S | SNN 109P | WT | SNN 109S | SNN 109P | WT | SNN 109S | SNN 109P | WT (at 5.46 Å) | SNN1 09S (at 5.62 Å) | SNN1 09S (at 7.00 Å) |
| -- | Oδ1(110)-OH(158) + Oδ2(110)-OH(158) | 1 | 0.97 | 0.64 | 1 | 0.64 | 0.46 | 0.69 | 0.58 | 0.36 | 0.25 | 0.95 | 0.51 | 0.04 |
| α-turn | N(112)-O(108) | 1 | 0.91 | 0.15 | 0 | 0.85 | 0.04 | 0 | 0.16 | 0.04 | 0 | 0.8 | 0.56 | 0 |
|  | N(113)-O(109) | 0 | 0 | 0.04 | 0.09 | 0 | 0.04 | 0 | 0.03 | 0.01 | 0 | 0 | 0.04 | 0 |
| β-turn | N(111)-O(108) | 0 | 0 | 0 | 0 | 0 | 0 | 0 | 0 | 0.01 | 0 | 0 | 0.01 | 0 |
|  | N(112)-O(109) | 0 | 0 | 0.45 | 0.73 | 0 | 0.63 | 0.02 | 0.55 | 0.08 | 0 | 0 | 0.08 | 0 |
| Back bone | distal β-sheet | 7 | 6.32 | 6.1 | 5.94 | 6.19 | 2.62 | 1.45 | 1.85 | 0.67 | 0.35 | 6.15 | 5.8 | 5.41 |

### Succinimide induces formation of conformational-lock, limiting loop flexibility

Molecular dynamics simulations (MDS) provide an opportunity to examine the impact of side-chain-backbone cyclization on the conformational flexibility of loop containing the succinimide modification. Experimental studies have established that the succinimide lacking MjGATase mutants SNN109S(*10*) and SNN109P are much less stable than the WT enzyme, precipitating at significantly lower temperatures (Fig. 2b). Diffraction quality crystals could not be obtained for the SNN109S mutant despite considerable effort. The structure of SNN109S was therefore modelled in PyMol(*48*) using the MjGATase structure as a template and energy minimized. The WT, SNN109S and SNN109P mutant structures were then subjected to REST2 MD simulations at a temperature range of 47-327 °C, followed by normal MD at temperatures of 27, 70 and 87 °C (see ‘Computational Methods’ section in the Supplementary Information for details including the necessity of REST2 that provides enhanced sampling over normal MD). In the WT enzyme, a strong 13-atom (C_13_, 5→1) hydrogen bond observed between Glu108 CO and Phe112 NH is maintained at temperatures up to 70 °C, indicative of the role of succinimide in stabilizing the α-turn motif. Interestingly, at 87 °C, there is a local conformational transition with population of Type II’ β-turn structures, stabilized by a 10-atom (C_10_, 4→1) hydrogen bond between the backbone CO of the succinimide ring and the Phe112 NH groups (Fig. 4a and Table 2). At 87 °C, both Asp110 and Leu111 populate a second distinct conformational state, corresponding to the formation of a type II’β-turn centred at residues 110-111 (Fig. 4d). The succinimide containing segment is thus conformationally constrained, restricting loop flexibility, even at the highest temperature examined through MDS. In the mutant SNN109S, the dominant conformations of the loop segment involve a β-turn H-bond between S109 CO and Phe112 NH groups. This interaction is almost completely lost at 87 °C (Fig. 4b and Table 2). We refer to these hydrogen bonds resulting in α-turn and β-turn motifs as the ‘conformational-lock’. In SNN109P mutant as well, the dominant conformation at 27 °C involves β-turn hydrogen bonds between P109 CO and Phe112 NH and D110 CO and K113 NH groups and these interactions are completely lost at higher temperatures (Fig. 4c and Table 2).

**Figure 4:**
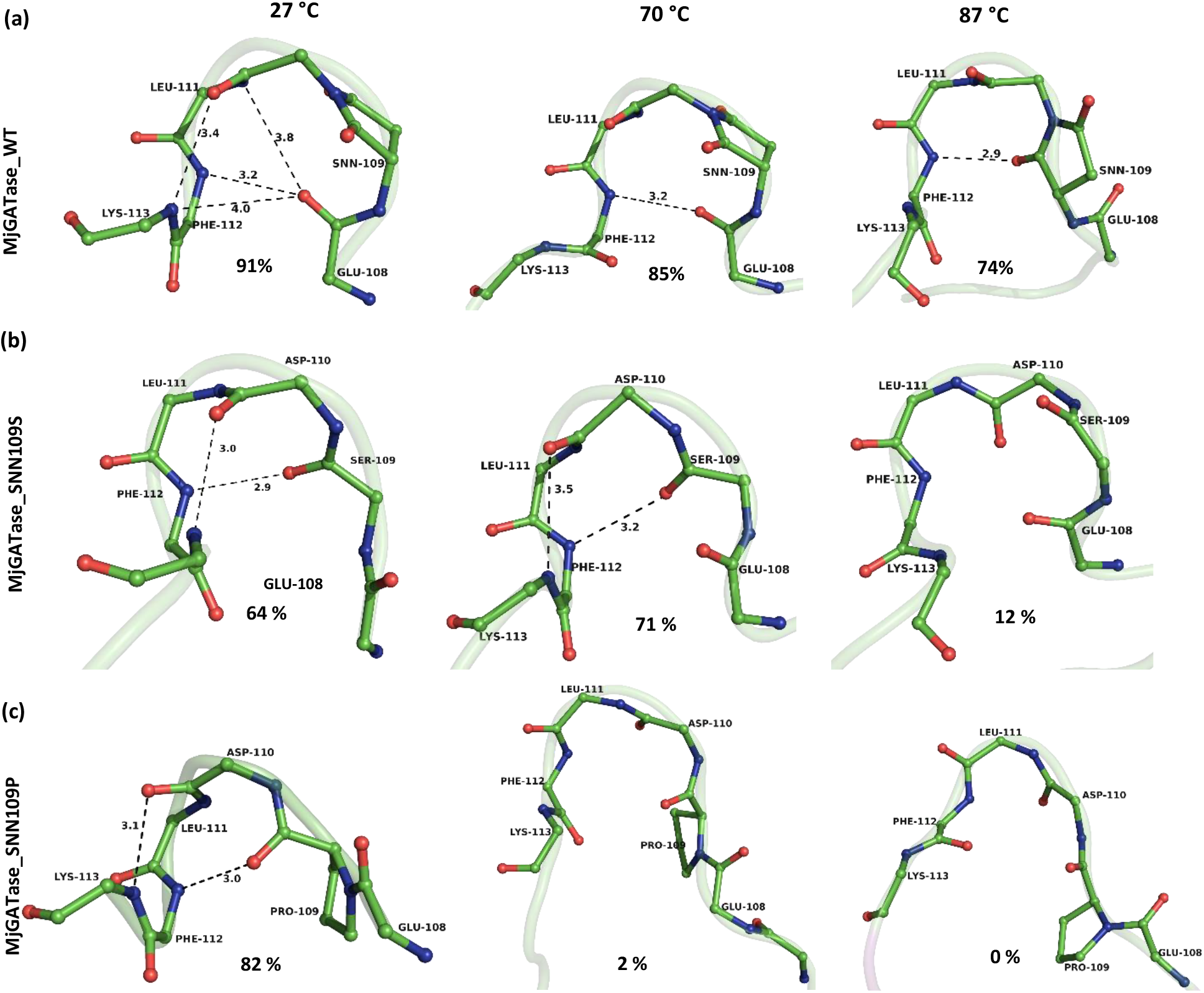
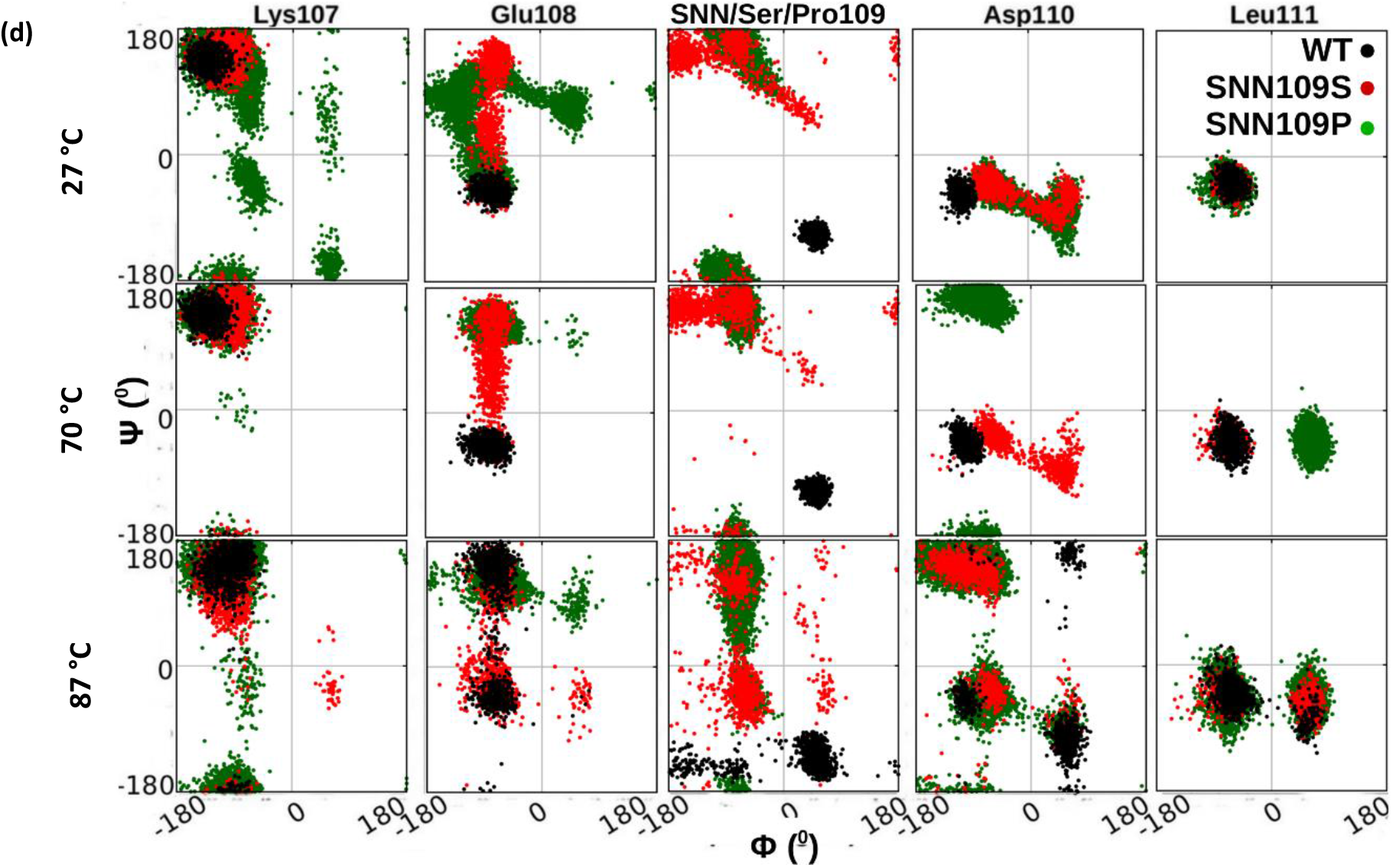
The contributions of *conformational-lock* as seen from simulations. (a) MD simulations suggest that the *conformational-lock* formed by the cyclization of residue at position 109 in MjGATase_WT is maintained at all temperatures although in different forms. β-turn exists in the structure at 87 °C. (b) The H-bonds that form α and β turns in the WT protein are replaced by a new set of H-bonds in the SNN109S mutant. At 27 °C, the H-bonds between Ser109 CO---Phe112 NH and Asp110 CO---Lys113 NH constitute two β-turn structural motifs. However, these bonds are weakened at 70 °C and are lost at 87 °C. (c) For SNN109P mutant, the H-bond between Pro109 CO---Phe112 NH constitutes a β-turn structural motif at 27 °C, which disappears at higher temperatures. The percentile strength of the conformational-lock measured as the probability of its occurrence in the corresponding simulation (Table 2) is mentioned in panels a, b and c. (d) The Φ, Ψ values (at three different temperatures) of residues around succinimidyl (WT), seryl (MjGATase_SNN109S) and prolyl (MjGATase_SNN109P) residues are plotted on Ramachandran map for comparison. Supplementary figure 8 provides a comparison of the degree of dynamics with respect to crystal structure for residues 107-111 in WT, SNN109S and SNN109P mutants at 27 °C.

Figure 4d compares the effect of temperature on the backbone torsion angles of the residues in the pentapeptide segment of the succinimide containing loop, residues 107-111, for the WT enzyme and the mutants. It is evident that in case of SNN109S and SNN109P, even at 70 °C, the Glu108, Ser109/Pro109 and Asp110 show considerable conformational excursions compared to the WT enzyme, as seen from the Ramachandran plots. It is interesting to note that the stability of the folded form of the loop consisting of residues 105-117 is considerably diminished, when the backbone conformational constraint at position 109 is through a prolyl residue vis-à-vis that achieved via the formation of a succinimidyl residue. The role of the succinimide induced “conformational-lock” was further probed using well-tempered metadynamics (WT-MTD) simulations at 27 °C, by applying a biasing force on the radius of gyration (Rg) of residues in the segment 108-115, containing the succinimide. This procedure promotes the breaking/forming of the “conformational-lock” several times, thereby boosting the sampling of conformational space of this region of the polypeptide chain to an extent greater than that through the REST2 protocol. However, this calculation has not been performed for the SNN109P mutant as the complete loss of conformational-lock at higher temperatures suggests an efficient conformational sampling for this system through REST2 itself (Table 2). Free energy profiles obtained from WT-MTD simulations show that the absence of the succinimide constraint in the SNN109S mutant allows this loop to exist in a locally, unfolded conformation, at a higher free energy (at the minimum corresponding to Rg value ∼ 7 Å). In contrast, there is little effect on the local conformation of this segment in the WT enzyme (Fig. 5a, Supplementary Movie 2). The free energy profiles in Figure 5a establish two distinct minima in the SNN109S mutant corresponding to the folded and less ordered conformations of the loop (Supplementary Movie 3). This observation is further supported by gas phase folding simulations of the unfolded 105-117 peptide segment with SNN at 109 position, wherein it is seen to fold in a more compact manner with a β-turn H-bond compared to that in the absence of succinimide (Supplementary Figure 6). Additionally, the MD simulations reveal that succinimide formation also has a long-range effect that limits the conformational lability of segments of the polypeptide chain distal to the site of modification.

**Figure 5:**
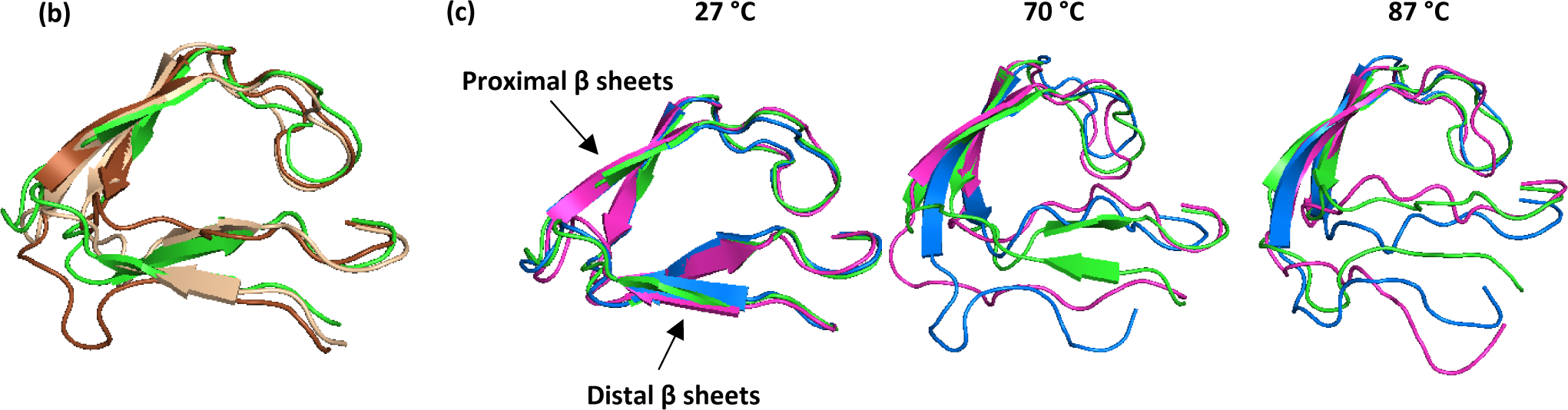
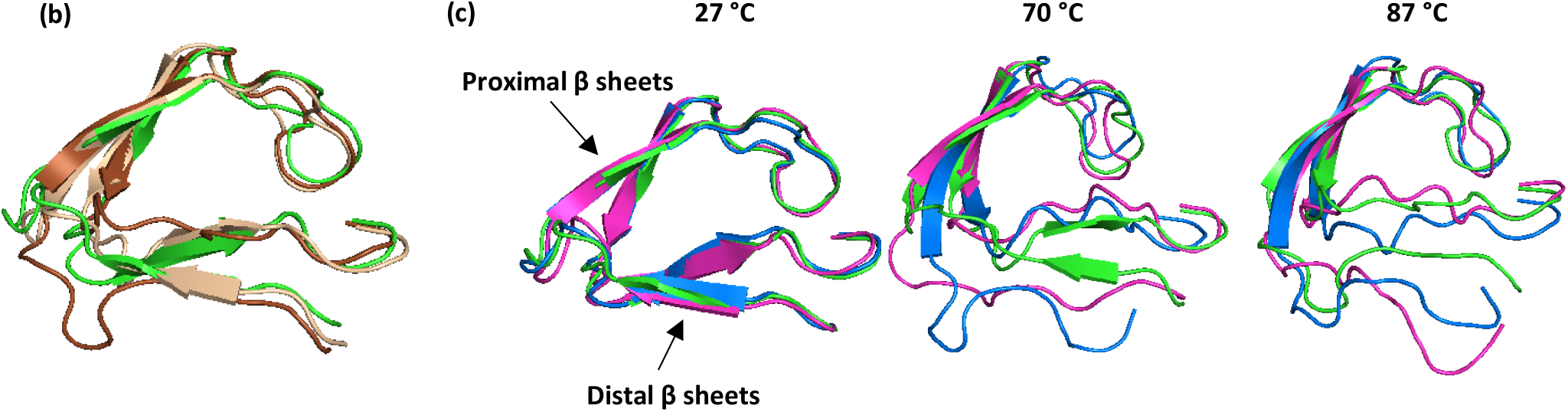
Short-range and long-range impacts of SNN on MjGATase structure, unravelled by simulations. (a) Free energy profiles with respect to the radius of gyration of residues near SNN (residues 108-115) calculated from metadynamics simulations. Conformation at each of the minimum are shown. Free energy minimum conformation of WT (green) shows a significantly good overlap with crystal structure (cyan). The SNN109S mutant can cross the free energy barrier to exist in a higher energy state in a completely unfolded conformation. (b) Overlay of the extended β-hairpin region of WT at 27 (green), 70 (pale brown) and 87 °C (brown). (c) Overlay of the extended β-hairpin region of WT (green), MjGATase_SNN109S (blue) and MjGATase_SNN109P (magenta) mutants.

Figures 5b and 5c show the effect of increasing temperature on the polypeptide chain spanning residues 86-136. The hairpin formed by the flanking antiparallel β-strand segments residues 97-105 and 115-124, connected by the loop 106-114, is further connected by three and two residue loops, respectively, to a second antiparallel β-sheet segment encompassing the β-strands spanning residues 86-93 and 127-136 (distal β-sheets). The disruption of the antiparallel β-sheet distal to the succinimide loop segment is evident in the SNN109S and SNN109P mutants at 70 and 87 °C (Fig. 5c and Table 2). In contrast, the overall fold of the polypeptide chain is maintained at high temperatures in the WT enzyme (Fig. 5b, c and Table 2, Supplementary Movies 4-9). A similar trend is reflected in the root mean square fluctuation (RMSF) of residues and fraction of native contacts with respect to crystal structure obtained from simulations (Supplementary Figure 7). Inspection of Figure 1c suggests that long range tertiary contacts involving the SNN109-Phe112 segment with Leu139, Lys151, Tyr158 and Phe162 residues from the core β sheets may be the contributing factors in the transmission of the structure stabilizing effect of the succinimide (Supplementary Table 1).

## Conclusions

Proteins from thermophiles have been shaped by the selective pressures of evolution to acquire structural features that determine their stability at temperatures as high as 80-120 °C. Despite the accumulation of a large body of structural and mutational data on proteins from hyperthermophiles, unambiguous correlates between sequence and conformational motifs contributing to thermal stability remain elusive(*49*–*51*). In the case of MjGATase being analysed in this report, succinimide formation by side-chain backbone cyclization, in an exposed loop segment contributes to hyper-thermostability. In the absence of the backbone constraint, the SNN109S mutant is stable up to a temperature of 85 °C. Locking residue 109 into the five membered succinimide ring freezes the backbone torsion angle Ψ to a value of 42.6°, placing it in the bottom right quadrant of the Ramachandran Map; a region of conformational space energetically unfavourable for L-amino acid residues in proteins. The promotion of intramolecular backbone hydrogen bonds at the loop tip (residues 106-114) is clearly a factor limiting loop dynamics as evidenced from the REST2 and WT-MTD simulations. Additionally, an important tertiary non-bonded contact is made between the Leu139 side-chain methyl group and the CβH2 of the Asn 109 residue which is now rigidly held within the succinimide ring. Long range tertiary contacts involving the SNN109-Phe112 segment with Leu139, Lys151, Y158 and F162 residues serves to transmit the structural stabilization, imposed by post-translational succinimide formation, to the distal active site segment. In proteins, the L-Pro residue is the sole amino acid in which a backbone cyclic constraint limits the N-C^α^ torsion angle (φ) to values close to −60°, a feature which promotes the formation of locally folded β-turn conformations(*52*). The SNN109P mutant harbouring a prolyl residue in place of succinimide, generated in this study, showed remarkably lower thermal stability than the WT enzyme. Although the segment corresponding to SNN loop in the crystal structure of the mutant displayed the presence of two consecutive β-turns; at higher temperatures, it was unable to retain the conformational-lock stabilizing the peptide segment in a folded state, as evidenced from the MD simulations. Further, many contacts of the SNN loop with the core of the protein that play a key structure stabilising role in the WT are absent in MjGATase_SNN109P. This indicates that although proline can decrease the conformational entropy of unfolding(*53*), this entropic contribution is masked by decreased enthalpic contributions as a consequence of loss of van der Waals and electrostatic interactions. Together, these results emphasize that the presence of a succinimidyl modification provides a superior constraint both entropically and enthalpically and that the structure stabilizing role of a specific amino acid residue is dependent on the location of that residue and its environment in the protein structure. Structural studies of model peptides reveal that β II’ conformations are favoured in L-Asu peptides, with the Asu residue occupying the *i+1* position of the turn motif(*54, 55*). In the MjGATase structure, the Asu (SNN)109 residue occupies the *i+1* position of an α-turn, a motif formed by the local conformations adopted by the succeeding residues Asp110 and Leu111. The recent demonstration of the effectiveness of a succinimidyl peptide as an inhibitor of Taspase, a leukaemia related threonine protease(*56*) may stimulate further conformational analysis of succinimide containing peptides. A very recent publication has also demonstrated the structure stabilizing effect of an iso-aspartyl residue in MurA from *Enterobacter cloacae*, highlighting the emerging role of backbone protein modifications(*57, 58*). As mentioned earlier, the observation of stable succinimides in proteins is uncommon and only a few examples are available in the Protein Data Bank(*5*–*8*). In MjGATase, the crystal structure and the MD simulations taken together establish that steric and electrostatic shielding by the Asp110 residue on one face of the succinimide ring, and by the backbone Glu108 CO (through potential n→π* interactions with carbonyl groups of succinimide) on the other face preclude approach of solvent water, thereby preventing hydrolysis. The hydrolytic instability of the D110G mutant demonstrated earlier(*10*) lends support to this interpretation. The key residues involved in interacting with the succinimide at position 109 are highly conserved in sequences of GATase from archaeal thermophiles, suggesting an evolutionary adaptation of sequence to meet the demands of thermal stability. The results presented above emphasise the utility of REST2 simulations in enhanced sampling of polypeptide conformational space, permitting examination of relatively minor structural changes on the unfolding of inherently stable proteins. A comparison of the simulations carried out on the WT enzyme and the mutants, SNN109S and SNN109P reveal complete unfolding of the distal β-sheet formed by antiparallel strands, 86-93 and 127-136 in the mutant proteins. Interestingly, in the structure of MjGATase_SNN109P, the electron density corresponding to residues 93-95 (the loop region connecting distal β-sheets) is missing. Thus, both simulation and experiments suggest that the effects of changes in the SNN-harbouring loop segment are not localized but are cascaded further into the protein. In contrast, in the WT enzyme, minor conformational changes are observed at 87 °C but the integrity of the structure over the core region 86-136 is maintained, affirming the role of the succinimide. As a caveat, it should be noted that although REST2 and WT-MTD simulations offer enhanced conformational sampling, a detailed view of the complete pathway of unfolding is beyond the scope of the current simulations. Deamidation of Asn residues is a commonly encountered phenomenon contributing to “wear and tear” of proteins(*59*). The facile hydrolysis of the succinimide intermediate can lead to Asp/isoAsp formation and stereochemical inversion, modifications that can be detrimental to activity(*1, 3, 4*). The MjGATase example illustrates how protein sequences can adapt an apparently detrimental process to serve the function of enhancing thermal stability, under an evolutionary imperative.

## Methods

### Generation of SNN109P mutant of MjGATase

SNN109P mutant was generated using standard procedures as described in Supplementary methods and confirmed by DNA sequencing.

### Purification of MjGATase_WT and mutant proteins

The genes for WT and mutant enzymes were cloned in pST39 expression vector and proteins were expressed in Rosetta (DE3) pLysS *E. coli* overexpression strain. Proteins were purified following protocols similar to that used for MjGATase WT(*10*), described in detail in supplementary methods.

### Mass spectrometric analysis of MjGATase_SNN109P and its tryptic digest

Mass spectra of purified MjGATase_SNN109P were acquired using Q Exactive HF orbitrap mass spectrometer (Thermo Scientific, USA). Details of the method are provided in supplementary methods.

### Thermal precipitation of proteins

Qualitative evaluation of thermal stability of WT and mutants of MjGATase was done by heating the proteins at five different temperatures followed separation of the precipitate and examination of soluble fraction by SDS-PAGE. Details of the method are provided in supplementary methods.

### Circular Dichroism

CD was performed to compare structure and stability of WT and SNN109S with SNN109P following standard protocol detailed in supplementary methods.

### Crystallization, Data collection, structure solution and refinement

Crystallization of WT and mutants were set up under 1:1 mixture of silicon and paraffin oil using the micro-batch method(*60*) and crystals were obtained only at 4 °C. X-ray diffraction data on all crystals were collected using Rigaku RU200 X-ray diffractometer equipped with a rotating anode type light source with an osmic mirror that gives a monochromatic light source of wavelength 1.54179 Å. Diffraction images were processed using iMOSFLM(*61*). Structure solutions were obtained using molecular replacement (MR) method using Phaser module of CCP4(*62*). For WT crystal, structure of GATase from *Pyrococcus horikoshii* (PDB ID: 1WL8) was used as template. For refinement of the structures, Refmac(*63*) module of CCP4 and AutoBuild module of Phenix(*64*) were used. Details of crystallization, data collection, structure solution and refinement are provided in supplementary methods.

### Interaction potential (force field) of succinimidyl residue

The force field parameters of succinimidyl residue were extracted from existing database of structurally similar molecules and benchmarked against crystal structures as well as quantum chemical and classical mechanical simulations. Details of force field parameterization and developed force field parameters are provided in supplementary methods.

### Regular MD simulation

MD simulations for WT MjGATase, WT PhGATase and SNN109S, SNN109P and D110G mutants of MjGATase were performed at 27 °C for 300 ns each. Details of the simulation protocol are provided in supplementary methods.

### REST2 followed by regular MD simulation

Replica exchange with solute (enzyme) scaling (REST2)(*34*) simulations were performed separately for the WT MjGATase and its SNN109S and SNN109P mutants (starting from the last frame of the corresponding regular MD simulation) with 28 replicas for each system in the effective temperature range 47-327 °C for the tempered region (protein) while the solvent was still in reference temperature of 47 °C for all replicas. The replicas were allowed to exchange every 10 ps. The average exchange probability between the successive replicas was found to be 35%.

After 250 ns long REST2 simulation, the last frame of the fourth and sixth replicas (with REST2 “effective” temperatures of 70 and 87 °C, respectively) were warmed up gradually to 70 and 87 °C, respectively. Regular MD was then performed for 300 ns for these systems at these temperatures. The analyses presented here are from these 200-300 ns portion of the normal MD trajectories, post the REST2 run. All the figures presented as snapshots from MD simulation were taken from the conformation of the central member of the most predominant cluster obtained from cluster analysis. Details of the REST2 simulation protocol and validation are provided in supplementary methods.

### Well-tempered metadynamics (WT-MTD) simulations

The free energy profile for WT MjGATase and its SNN109S mutant with respect to the radius of gyration (collective variable or CV) of the residues from the succinimide containing loop (residues 108-115) was reconstructed from 2200 ns long WT-MTD(*35*) at 27 °C. Details of the WT-MTD simulation protocol and validations are provided in supplementary methods.

### Natural Bond Orbital (NBO) analysis

Energetics of the n→π* interactions and n→σ* interactions (hydrogen bond) contributing toward the stabilization of succinimidyl residue were obtained through NBO analysis(*44*) on quantum optimized (only electronic level, keeping the atomic position fixed) structure of the polypeptide sequence 108-112 extracted from the crystal structure of WT MjGATase. Details of this analysis are provided in supplementary methods.

### Structure analysis and figure rendering

Figures of three-dimensional structures were drawn with PyMol(*65*) (DeLano Scientific LLC, http://pymol.sourceforge.net/)

## Acknowledgements

We acknowledge the technical support from X-ray facility at Molecular Biophysics Unit, Indian Institute of Science, Bangalore, India where the crystals were diffracted. We acknowledge assistance from Dr. Udupi A. Ramagopal (Poornaprajna Institute of Scientific Research, Bangalore, India), Prof. M.R.N. Murthy, Prof. B. Gopal and Dr. K.V. Abhinav (Molecular Biophysics Unit, Indian Institute of Science, Bangalore, India) in aspects of crystal structure solution. We also acknowledge Mass Spectrometry facility at Molecular Biology and Genetics Unit, Jawaharlal Nehru Centre of Advanced Scientific Research, Bangalore and Ms. Sunita Prakash and mass spectrometry facility of Molecular Biophysics Unit, Indian Institute of Science, Bangalore, India for mass spectra. S.D. and S.B. thank CDAC’s NPSF on whose computing platform, a part of these calculations was performed. S.D., A.B. and S.K. acknowledge CSIR for JRF and SRF fellowships. A.D. acknowledges JNCASR institutional fellowship. A.C. acknowledges Department of Biotechnology for JRF and SRF fellowships. P. B is supported by the Department of Science and Technology under the Year of Science Chair Professorship scheme. For H.B., this research was funded by the Department of Biotechnology, Ministry of Science and Technology, Government of India Grants BT/PR11294/BRB/10/1291/2014, BT/PR13760/COE/34/42/2015, and BT/INF/22/SP27679/ 2018; Science and Engineering Research Board (SERB) CRG/2019/004150/IBS, Department of Science and Technology, Government of India Grant EMR/2014/001276; JC Bose Fellowship, SERB and institutional funding from the Jawaharlal Nehru Centre of Advanced Scientific Research, Department of Science and Technology, India.

## Author Contributions

AVD, SK purified protein, obtained crystals

AVD, AB, SK collected data, solved structures

AC generated mutant, purified protein, spectroscopic studies

SD, TK generated force field for succinimidyl residue; SD, SB carried all simulation studies

AVD, PB, HB analysed structures

AVD, SD, PB, SB, HB wrote the manuscript

## Competing interests

The authors declare no competing interests.

## Notes

### Competing Interest Statement

The authors have declared no competing interest.

### Summary of Updates

Table 1 revised; author order updated

