## Supplementary Table 1 for "Structural basis for the hyperthermostability of an archaeal glutaminase induced by post-translational succinimide formation"

**Supplementary Table 1**: Tertiary contacts (van der Waals contacts (vdW) and hydrogen bonds (H-bond) measured by the average minimum distance between non-hydrogen atoms of corresponding residue pairs from MD simulations. These network of tertiary contacts correlates distal β-sheet from extended β-hairpin and active site residues with the residues present in succinimide containing loop. For WT-MTD, the position of free energy minimum with respect to radius of gyration (Rg) of residues 108-115 are mentioned within parenthesis and the average minimum distances are calculated over configurations around the corresponding free energy minimum (for WT: 5.36-5.56 Å; for SNN109S mutant, lower free energy minimum: 5.52-5.72 Å; for SNN109S mutant, higher free energy minimum: 6.90-7.10 Å). X denotes SNN, Ser and Pro, respectively for WT, SNN109S and SNN109P. *For crystal structure, it is not an average number. The weakening of tertiary contact for SNN109S mutant compared to that of WT may not be found in WT-MTD for those contacts which does not involve residues within 108-115, as biasing force had been applied only to residues 108-115.

| Type | Residue pair(s) | Average* minimum distance (Å) | | | | | | | | | | | | | |
|---|---|---|---|---|---|---|---|---|---|---|---|---|---|---|---|
| | | Crystal* | | 27 ℃ | | | 70 ℃ | | | 87 ℃ | | | WT-MTD (27 ℃) | | |
| | | WT | SNN 109P | WT | SNN 109S | SNN 109P | WT | SNN 109S | SNN 109P | WT | SNN 109S | SNN 109P | WT (at 5.46 Å) | SNN 109S (at 5.62 Å) | SNN109S (at 7.00 Å) |
| vdW | X109-L139 | 3.81 | 3.57 | 3.5 | 3.72 | 3.93 | 3.47 | 3.81 | 4.02 | 4.13 | 5.17 | 4.34 | 3.49 | 5.96 (SNN got flipped) | 3.75 |
| vdW | X109-K151 | 3.5 | 5.79 | 4.15 | 4.67 | 5.06 | 3.82 | 5.71 | 6.4 | 6.15 | 7.22 | 6.8 | 3.97 | 6.03 | 6.34 |
| Hbond | D110-Y158 | 2.7 | 3.56 | 2.86 | 3.38 | 2.94 | 3.38 | 4.23 | 3.42 | 3.86 | 4.76 | 3.83 | 2.85 | 3.75 | 6.17 |
| vdW | L111-V160 | 3.79 | 4.32 | 4.32 | 4.54 | 4.6 | 4.38 | 5.02 | 8.82 | 4.55 | 5.36 | 8.02 | 4.34 | 4.98 | 6.29 |
| vdW | F112-F162 | 4.14 | 3.98 | 4.24 | 4.3 | 4.57 | 4 | 4.15 | 4.98 | 3.98 | 4.17 | 5.19 | 4.33 | 4.27 | 7.24 |
| Hbond | V160-C76 | 2.82 | 2.78 | 2.95 | 2.87 | 2.87 | 3.97 | 3.32 | 3.46 | 4 | 3.17 | 2.94 | 2.88 | 4.49 | 4.52 |
| vdW | L151-F136 | 3.18 | 3.22 | 3.46 | 3.42 | 3.42 | 3.46 | 3.52 | 3.42 | 3.44 | 3.54 | 3.48 | 3.42 | 3.42 | 3.44 |
| vdW | F136-Y86 | 3.58 | 3.69 | 3.78 | 3.74 | 3.8 | 3.95 | 6.73 | 4.11 | 3.83 | 5.75 | 6.7 | 3.78 | 3.88 | 3.84 |
