## Supplementary figures for "Structural basis for the hyperthermostability of an archaeal glutaminase induced by post-translational succinimide formation"

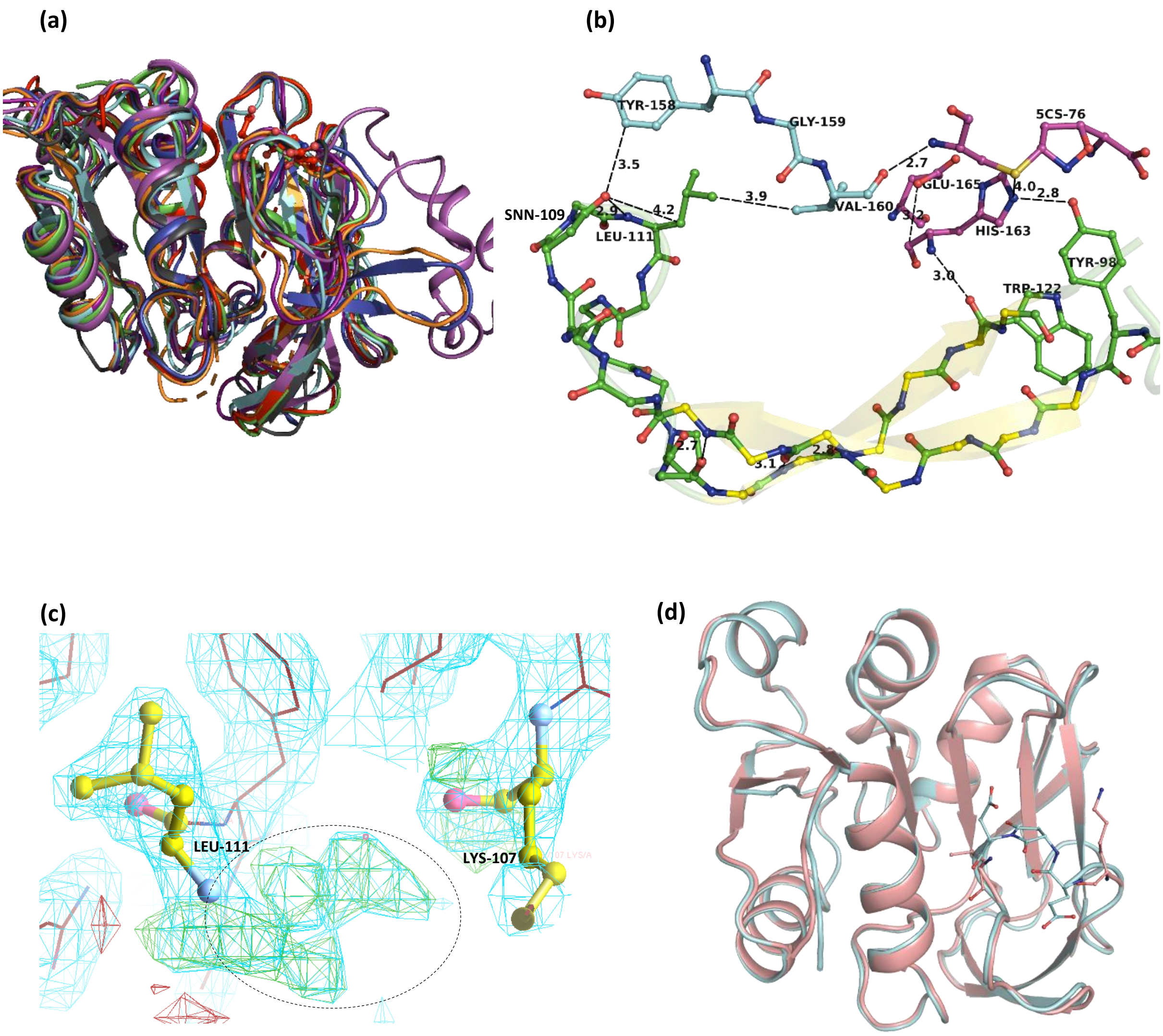


**Supplementary Figure 1: Structural features of apo, acivicin bound WT protein and MjGATase_D110G**
(a) Similarity of the overall folds in GATase subunits of GMP synthetase from eight different organisms. The central region comprising of seven mixed β-strands is conserved in all these enzymes. Structures of GATase subunit/domain of GMP synthetases deposited in PDB were superimposed using PyMol. *Methanocaldococcus jannaschii* (7D40), red; *Pyrococcus horikoshii* (1WL8), green; *Plasmodium falciparum* (4WIN), purple; *Escherichia coli* (1GPM), blue; *Homo sapiens* (2VPI), grey; *Coxiella burnetii* (3TQI), orange; *Neisseria gonorrhoeae* (5TW7), pink; *Thermus thermophilus* (2YWC), cyan. (b) Position of acivicin covalently-bound to Cys76 (5CS) with respect to succinimidyl containing β-hairpin. Active site residues are coloured in magenta, β-hairpin residues in green and the connecting residues in cyan. (c) Fragmented electron density in the region 108-110 in MjGATase_D110G (blob of fragmented density is circled). (d) Superimposition of the overall structure of WT (cyan) with the structure of MjGATase_D110G (pink).

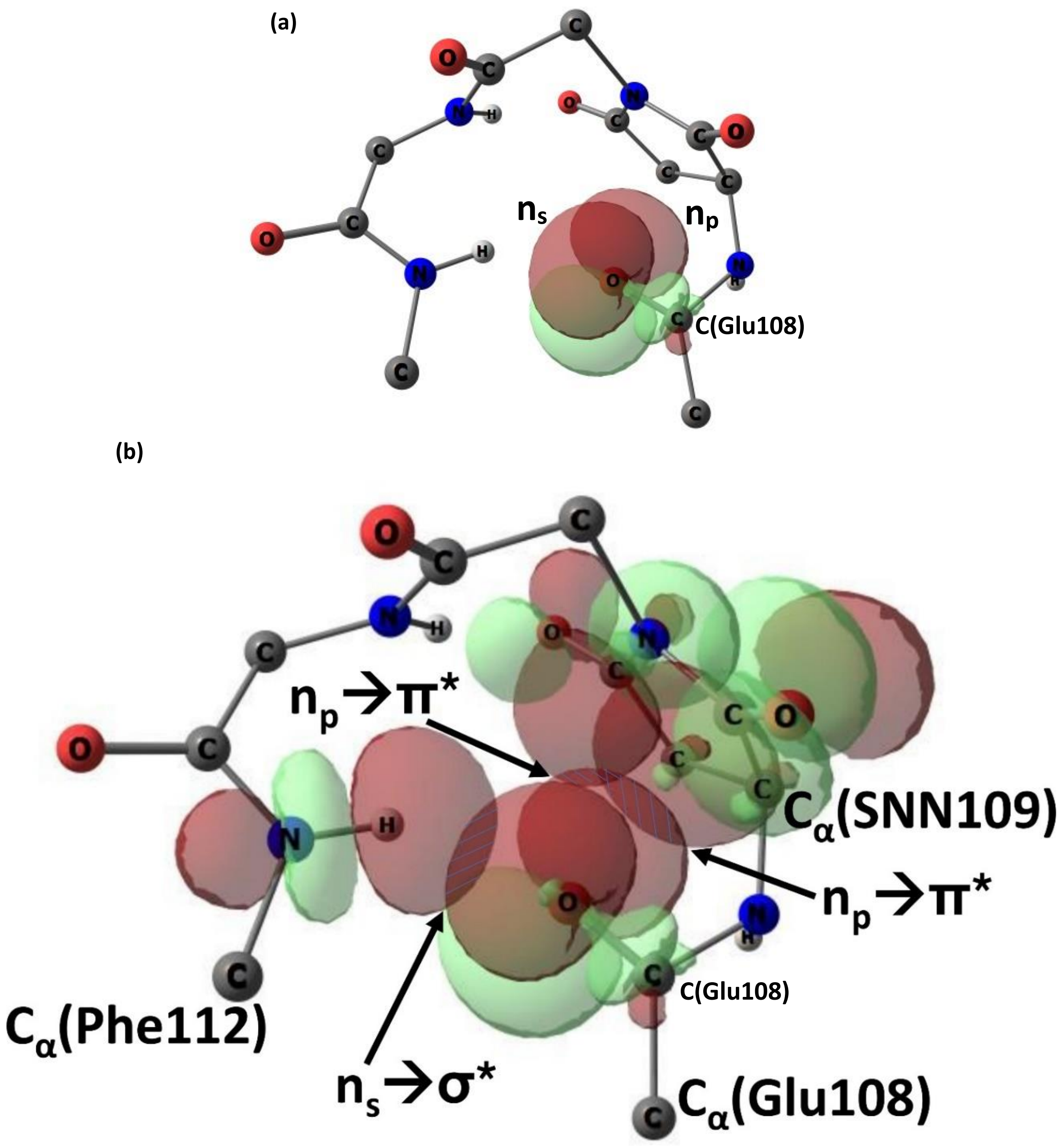


**Supplementary Figure 2: Interplay between hydrogen bond (n→σ* interaction) and n→π* interaction.** (a) De-mixing of two $sp^2$-type orbitals of backbone carbonyl oxygen of Glu108 into one s-type (containing lone pair 1 or LP1) and one p-type (containing lone pair 2 or LP2) orbitals due to the change of the C=O…HN angle (changes from ~$120^0$ to ~$160^0$) of the local α-turn hydrogen bond between Glu108 CO and Phe112 NH groups imposed by the global fold of peptide segment 108-112 induced by succinimide. A value of ~$91^0$ of the angle C=O(108)...C(109) in the crystal structure suggests almost complete de-mixing of these lone pair orbitals. (b) This de-mixing allows facile delocalization of lone pair LP2 from the p-type orbital of backbone carbonyl oxygen of Glu108 to the π* orbital of backbone and side-chain carbonyl groups of the following succinimidyl residue. Whereas, the lone pairs LP1 residing on s-type orbital of backbone carbonyl oxygen of Glu108 delocalized mostly to the σ* orbital of the backbone NH group of Phe112 forming an α-turn hydrogen bond. The natural bond orbital (NBO) analysis ( https://nbo6.chem.wisc.edu/) results in the stabilization energies as follows: for n→π* (with backbone CO of succinimide) 2.61 kcal/mol (17.33% LP1, 88.02% LP2), for n→π* (with side-chain $C_γO_γ$ of succinimide) 0.19 kcal/mol (2.17% LP1, 5.37% LP2), for n→σ* (hydrogen bond) 2.39 kcal/mol (80.50% LP1, 6.61% LP2). Except succinimide residue, only backbone atoms (NH-$C_α$-CO) for rest of the residues are shown for clarity. Figures are made with ChemCraft ( https://www.chemcraftprog.com)

.

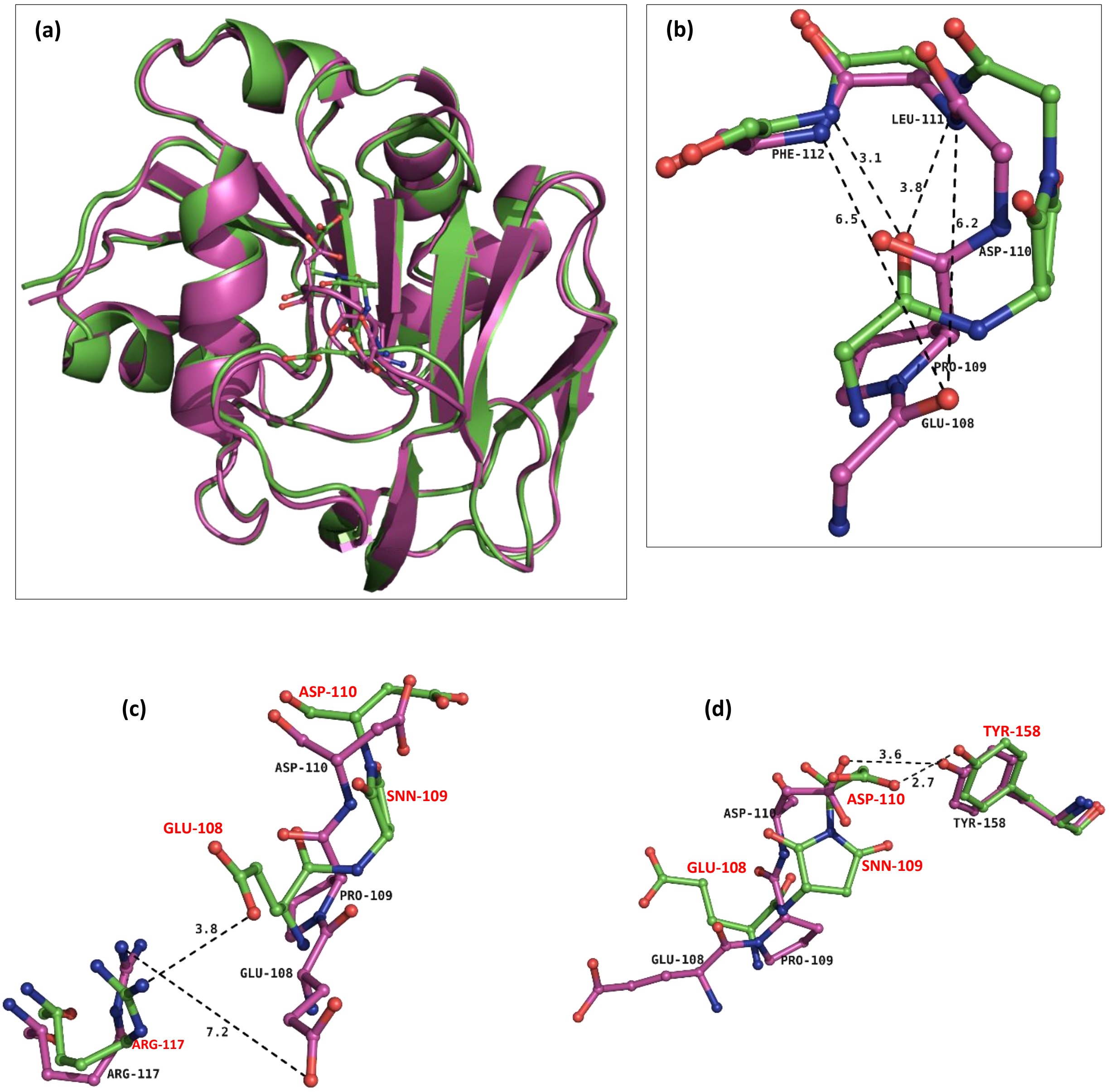


**Supplementary Figure 3: Comparison of the structures of MjGATase_WT and MjGATase_SNN109P.** (a) Superposition of the crystal structures of MjGATase_WT and MjGATase_SNN109P. (b) Zoomed in view of the superposition of the tripeptide region 108-110 of MjGATase_WT and SNN109P mutant shows that the backbone carbonyl group of Glu108 that is involved in formation of α-turn and β-turn structural motifs in WT protein is moved away in SNN109P structure and is therefore, no longer able to form the corresponding H-bonds. (c) Salt bridge between side-chains of E108 and R117 that is seen in WT is lost in the SNN109P mutant due to shift in orientation of E108 residue. (d) Superposition of WT and SNN109P mutant structures illustrating change in interaction between Asp110 and Tyr158.

**(a)**

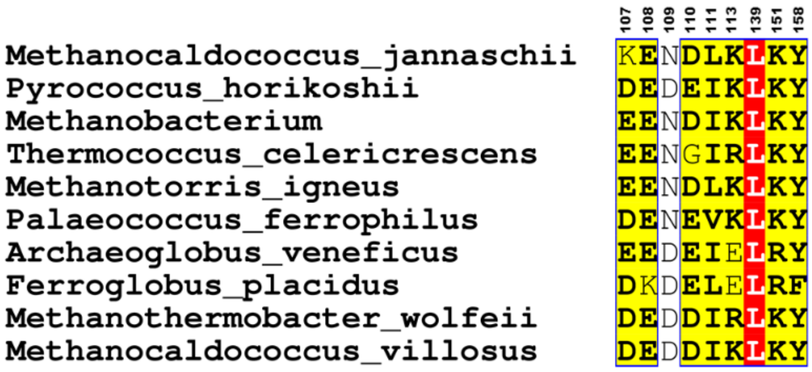
107 108 109 110 111 113 139 151 158
Methanocaldococcus_jannaschii KENDLKLKY
Pyrococcus_horikoshii DEDEIKLKY
Methanobacterium EENDIKLKY
Thermococcus_celericrescens EENGIRLKY
Methanotorris_igneus EENDLKLKY
Palaeococcus_ferrophilus DENEVKLKY
Archaeoglobus_veneficus EEDEIELRY
Ferroglobus_placidus DKDELELRF
Methanothermobacter_wolfeii DEDDIRLKY
Methanocaldococcus_villosus DEDDIKLKY

**(b)**

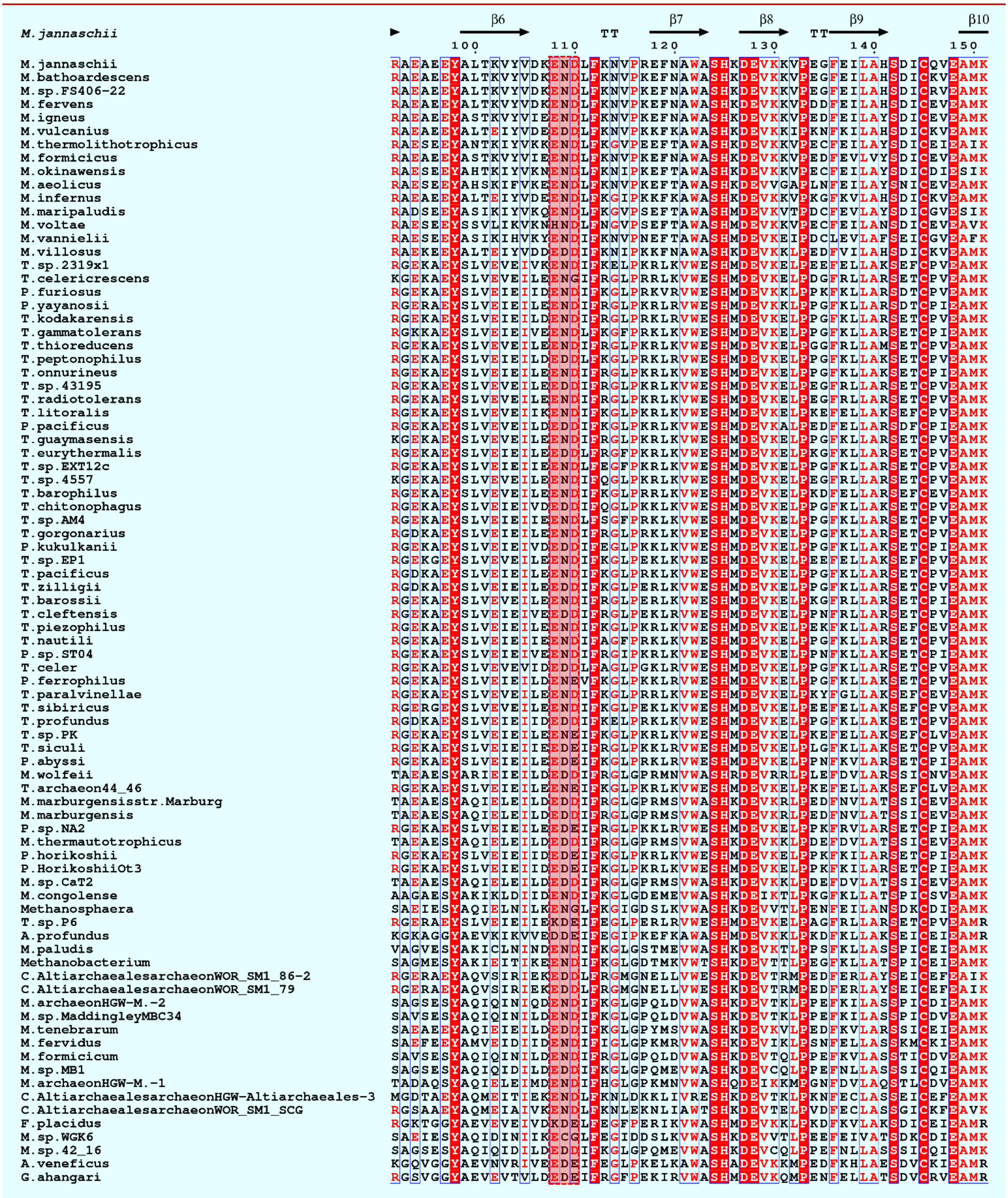


**Supplementary figure 4: Alignment of protein sequences of GATase subunit of GMP synthetases from archaea.** (a) Sequence conservation of residues in the vicinity of SNN across 10 archaeal GATases. (b) The protein BLAST performed using MjGATase as query sequence provided 100 hits which were curated to remove redundant sequences and the resultant 84 sequences were aligned using Clustal Omega. Aligned sequences were analysed and represented using ESPript 3.0. The conserved E(N/D)(D/E) sequence is highlighted in pink box.

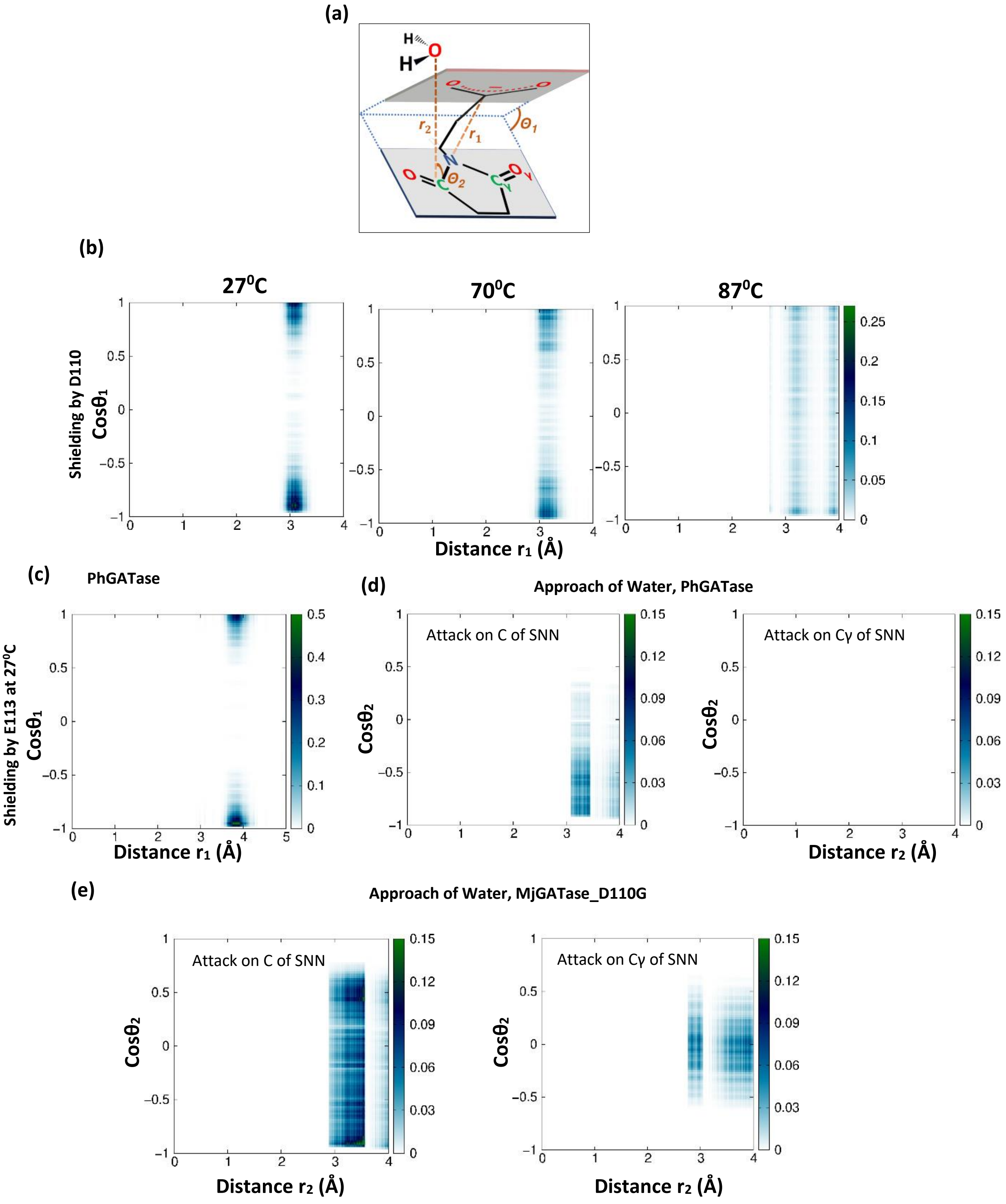


**Supplementary figure 5: Succinimide is shielded from hydrolysis by succeeding residue.** (a) Schematic representation of the distances and angles used in rest of the panels. See Figure 3 legend in the main text for details. (b) Probability distributions P($r_1$, cos$\theta_1$) are shown. $r_1$ represents N(SNN109)-C$_\gamma$(Asp110) distance and $\theta_1$ is the angle between the SNN-plane and side-chain carboxylate plane of Asp110. Values of $r_1$ and cos$\theta_1$ in the crystal structure are 2.93 Å and +0.96, respectively. (c) Similar results are obtained for *Pyrococcus horikoshii* GATase wherein at 27 $^0$C simulation, Glu113 shields SNN from hydrolysis as seen from probability distribution plot. (d) Probability distributions P($r_2$, cos$\theta_2$) showing the approach of water towards C and Cγ atoms of SNN in PhGATase. (e) Probability distributions P($r_2$, cos$\theta_2$) showing the approach of water towards C and Cγ atoms of SNN in MjGATase_D110G.

**(a)**

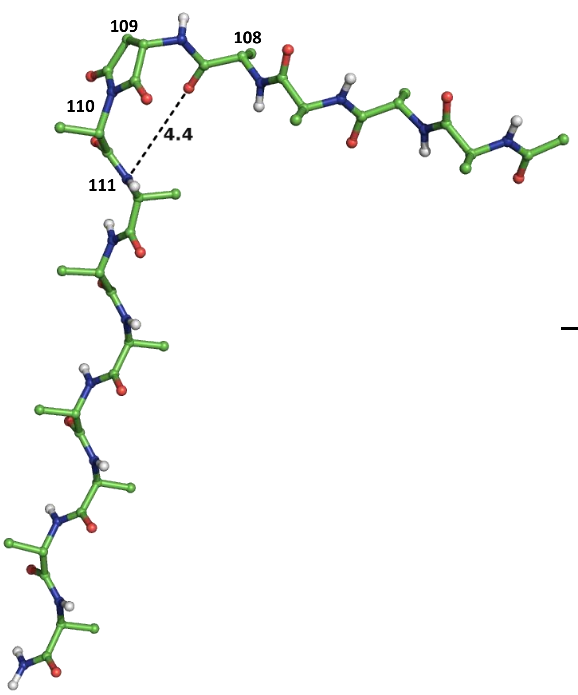
109
108
110
4.4
111
110
109
111
3.2
108
2.9
2.9
2.9

**(b)**

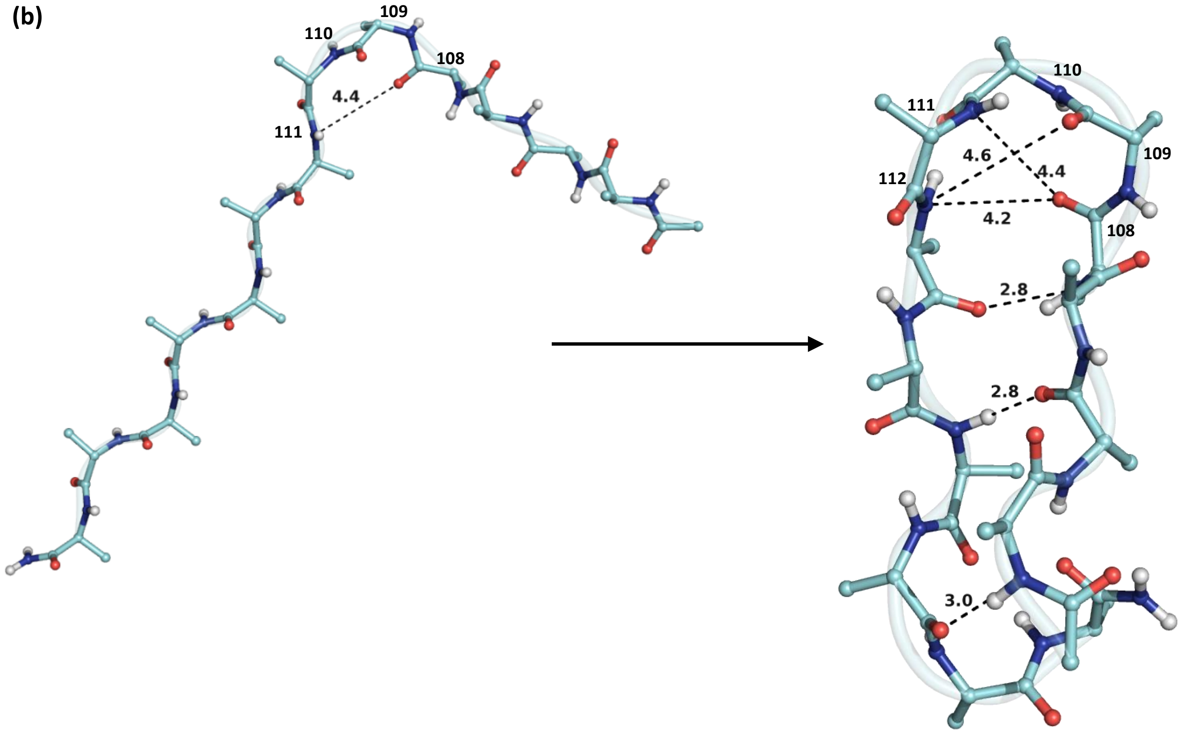
109
110
108
4.4
111
110
111
109
4.6
4.4
112
4.2
108
2.8
2.8
3.0

(c)

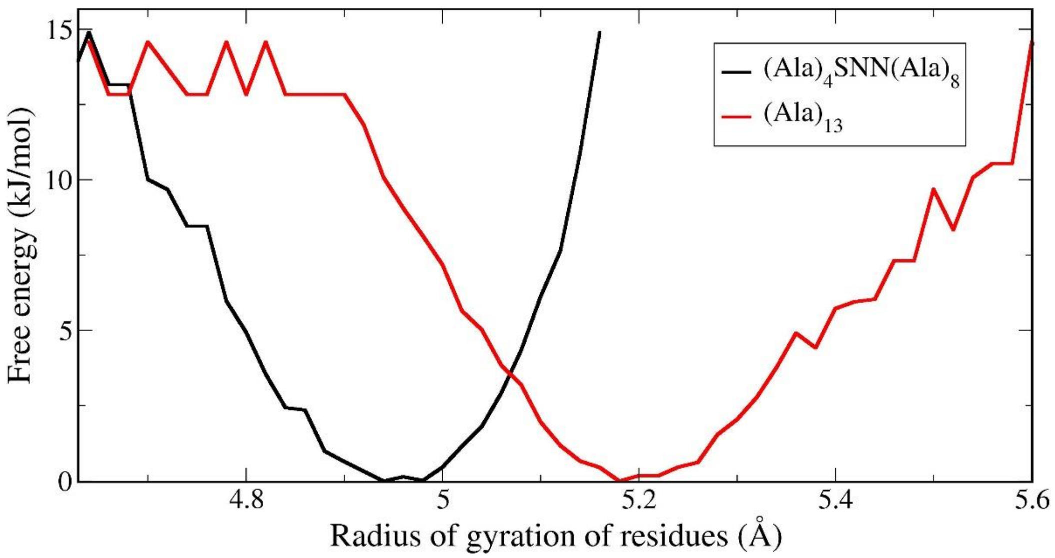


**Supplementary figure 6: MD simulations (for 100 ns at 27 $^{0}$C) of gas phase folding of succinimide containing poly Ala peptide.** (a) The peptide segment corresponding to residues 108-115 was taken from MjGATase crystal structure, followed by mutation of all the residues barring succinimide, to alanine. The N and C-terminals were then acylated and amidated, respectively. To get the unfolded structure, the Ramachandran dihedral angle pair for each residue except succinimide was kept at (-120$^{0}$, 120$^{0}$), whereas the values for the succinimidyl residue were taken from the crystal structure. MD simulation protocols were the same as mentioned in the Method section. MD simulation folds this peptide segment to form a β-turn hydrogen bond between the CO of residue 108 and N of residue 111. (b) Started from the same unfolded structure as 'a', but mutating succinimide as well to alanine. Although the polypeptide segment folds into a hairpin structure, owing to absence of succinimide, neither α nor β turn are formed. (c) Free energy profile with respect to the radius of gyration of residues calculated over 30-100 ns of the corresponding simulation trajectories of the peptide in gas phase shows that in the presence of succinimide the peptide segment folds in a more compact manner (by forming β-turn hydrogen bond). These profiles were obtained by Boltzmann inversion of the populations. The residues included for the calculation of radius of gyration of the peptide with SNN were Ala-SNN-$(Ala)_6$ with the control being $(Ala)_8$.

(a)

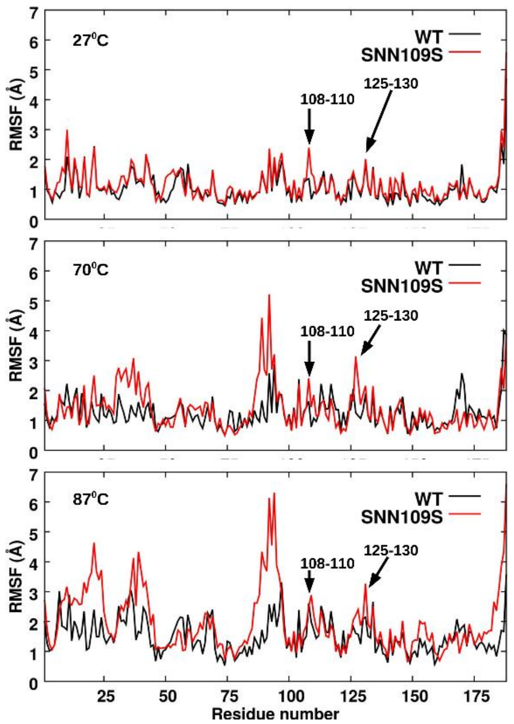


(b)

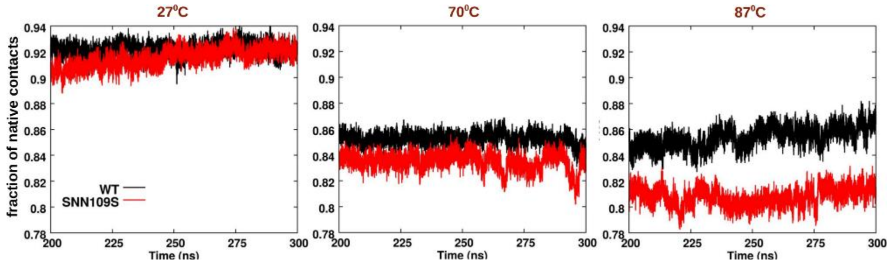


**Supplementary figure 7: Effect of conformational-lock on the entire enzyme elucidated by simulations.** (a) Comparison of root mean square fluctuation (RMSF) of residues between WT and MjGATase_SNN109S. (b) Comparison of the fraction of native contacts with respect to the crystal structure between WT and MjGATase_SNN109S.

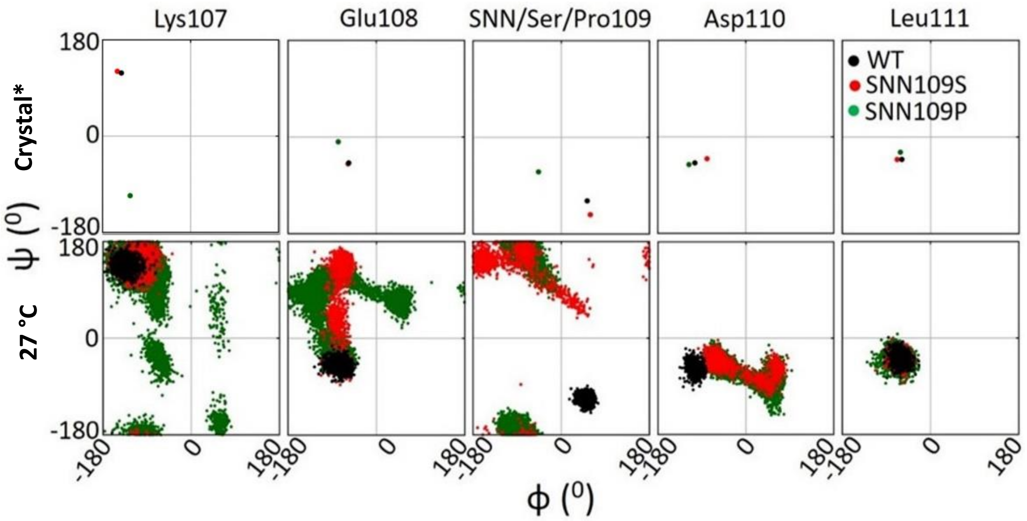


**Supplementary figure 8:** Ramachandran plot for residues near SNN109 (residues 107-111) in the crystal structure of WT and MjGATase_SNN109P. *The initial values for MjGATase_SNN109S were taken from structure generated (by mutating SNN109 of WT MjGATase to Ser) and energy minimized *in-silico*. (top panel). Bottom panel shows MD trajectory at 27 ℃ within the time frame of 200-300 ns in MjGATase_WT, MjGATase_SNN109S and MjGATase_SNN109P. The comparison of Ramachandran plot for any one of the residues between these two panels reveals that in the course of the MD simulation at 27 ℃, SNN109 in WT more or less retains its native conformation in Ramachandran torsional angle space, whereas in SNN109S and SNN109P mutants, Ser109 and Pro109, respectively, deviate from their initial conformations. These deviations had started at a very early stage of the MD production run.
