## Supplementary Information for "Structural basis for the hyperthermostability of an archaeal glutaminase induced by post-translational succinimide formation"

| **Author name** | **Email address** | **ORCID ID** |
|---|---|---|
| Aparna Vilas Dongre | | https://orcid.org/0000-0002-9124-7863 |
| Sudip Das | | https://orcid.org/0000-0001-8776-449X |
| Sundaram Balasubramanian | | http://orcid.org/0000-0002-3355-6764 |
| Tarak Karmakar | | https://orcid.org/0000-0002-8721-6247 |
| Anusha Chandrashekarmath | | https://orcid.org/0000-0003-1609-9490 |
| Asutosh Bellur | | https://orcid.org/0000-0002-5366-0630 |
| Padmanabhan Balaram | | https://orcid.org/0000-0002-6577-933X |
| Hemalatha Balaram | | https://orcid.org/0000-0002-8821-1288 |
| Sanjeev Kumar | | https://orcid.org/0000-0001-6653-0467 |

### Supplementary Information

#### Outline:

1) Supplementary Methods
   A) Experimental methods
   B) Computational methods
      i) Validation of computational methods
      ii) Force field parameters for succinimide

#### A) Experimental Methods

Phusion DNA polymerase, deoxynucleotide triphosphates (dNTPs) were procured from Thermo Scientific (USA). Primers were custom synthesized at Sigma Aldrich Co., India. All chemicals used were of highest purity and were procured from Merck & Co. (Sigma-Aldrich) Akta Basic HPLC, Q-sepharose resin were from GE healthcare life sciences, UK. Mass spectral studies were done at Molecular Biophysics Unit, Indian Institute of Science, Bangalore, India using a high resolution ESI-Q-TOF mass spectrometer (Maxis Impact, Bruker Daltonics, Germany), coupled to Agilent HPLC system and JNCASR mass spectrometry facility using Q Exactive$^{TM}$ HF (Thermo Scientific) mass spectrometer coupled to Easy nanoLC 1200 (Thermo Scientific). X-ray diffraction data for WT, D110G and SNN109P mutants were collected on Rigaku RU200 X-ray diffractometer equipped with a rotating anode type light source with an osmic mirror that gives a monochromatic light source of wavelength 1.54179 Å. An image plate of type MAR scanner, 345 nm was used for detection.

##### Generation of SNN109P mutant of MjGATase

SNN109P mutant of MjGATase was generated by site-directed mutagenesis using pST39_WTMjGATase as template. The primer pair used was forward primer: AGGTCTATGTAGATAAAGAACCAGATTTATTTAAAAACGTTCC and reverse primer GGAACGTTTTTAAATAAATCTGGTTCTTTATCTACATAGACCT. Introduction of mutation was confirmed by DNA sequencing (Supplementary Figure 9 a) and mass spectral analysis of the intact protein as well as MS-MS analysis of the tryptic digest of MjGATase_SNN109P protein (Supplementary Figure 9 b, c).

##### Protein purification

The WT and mutant enzymes were expressed and purified following protocols similar to that used for MjGATase WT(1). Gene for MjGATase_WT and mutants was cloned in pST39 expression vector and expressed into Rosetta (DE3) pLysS *E. coli* overexpression strain. Cells containing the plasmid construct were grown at 37 °C in terrific broth containing 100 μg ml$^{-1}$ ampicillin and 34 μg ml$^{-1}$

chloramphenicol to an $OD_{600}$ value of 0.6. and thereafter the cells were induced with IPTG to a final concentration of 0.3 mM and grown for additional period of 3 hours. At the end of 3 hours, cell pellet was collected by centrifugation at 3400 g for 10 minutes at 4 °C, and re-suspended in 30 ml of lysis buffer (20 mM Tris-HCl, 10% glycerol, 0.1 mM EDTA, 1 mM PMSF, 2 mM DTT, pH 8.0) and lysed by sonication. The cell lysate was centrifuged at 24,000 g for 30 minutes at 4 °C to remove cell debris. The supernatant was heated at 70 °C for 30 minutes to denature and precipitate *E. coli* proteins. The denatured endogenous bacterial proteins were removed by centrifugation at 24,000 g for 30 minutes at 4 °C. In order to precipitate nucleic acids, supernatant was treated with 0.01 % of polyethyleneimine and centrifuged at 24,000 g for 30 minutes. The resulting supernatant was loaded onto a Q-sepharose anion-exchange column equilibrated with buffer A (20 mM Tris-HCl, 10% glycerol, 0.1 mM EDTA, 1 mM PMSF, 2 mM DTT, pH 8.0) at a flow rate of 3.0 ml $min^{-1}$. The column was washed with 50 ml of buffer A. The protein was eluted with a linear gradient of buffer B (Buffer A containing 1 M NaCl). The elution was monitored by measuring absorbance at 280 nm. The fractions containing protein were collected, and concentrated using centrifugal concentrators (Centricon, Millipore, USA, 10.0 kDa MWCO). The protein concentration was estimated by the method of Bradford using BSA (1 mg $ml^{-1}$) as a standard.

**Mass spectrometric analysis of MjGATase_SNN109P and its tryptic digest**

Mass spectrum of the purified enzyme diluted in water : methanol (50:50 with 0.1% formic acid) from a stock concentration of 40 mg $ml^{-1}$ to 0.1 mg $ml^{-1}$ was acquired by direct infusion using a Hamilton syringe in a Q Exactive HF orbitrap mass spectrometer (Thermo Scientific, USA) and data was acquired in the positive ion mode.

MjGATase_SNN109P was subjected to in-gel trypsin digestion to acquire MS/MS spectra of peptides in order to confirm the presence of proline residue at position 109. In-gel tryptic digestion was performed using previously published protocol(2). Briefly, 30 μg purified protein was run on a 12% SDS-PAGE gel and stained with Coomassie Brilliant Blue R-250. Gel piece containing protein band was excised, cut into small pieces and de-stained using 100 mM ammonium bicarbonate/acetonitrile (1:1, vol/vol). Reduction and alkylation of de-stained protein was done using 10 mM DTT and 55 mM iodoacetamide (in 100 mM ammonium bicarbonate) respectively followed by dehydration using 100 % acetonitrile. The gel pieces were rehydrated by adding sufficient volume of trypsin buffer followed by addition of 10-20 μL ammonium bicarbonate to keep the gel pieces wet during enzymatic digestion. Digested peptides were extracted using extraction buffer (1:2; 5% formic acid/acetonitrile). The extracts were vacuum dried and stored until LC-MS/MS analysis. For analysis, each sample was re-dissolved in 0.1% (vol/vol) formic acid (in UHPLC/MS grade water) vortexed and centrifuged. Supernatant was carefully transferred to a fresh LC-MS sample vial. Mass spectral analysis was carried out on the Q Exactive HF orbitrap mass spectrometer (Thermo Scientific, USA) equipped with Easy nanoLC 1200 (Thermo Scientific). Data was acquired in positive ion mode using HCD fragmentation method. Data was viewed and analysed using Qual Browser in Xcalibur and Proteome-Discoverer 2.3 (Thermo Scientific), respectively.

**(a)**

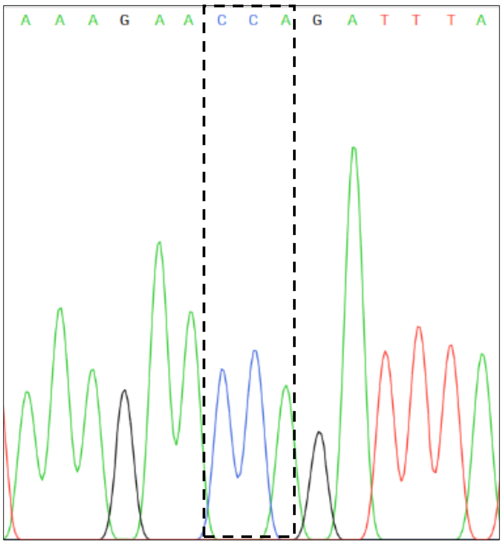

**(b)**

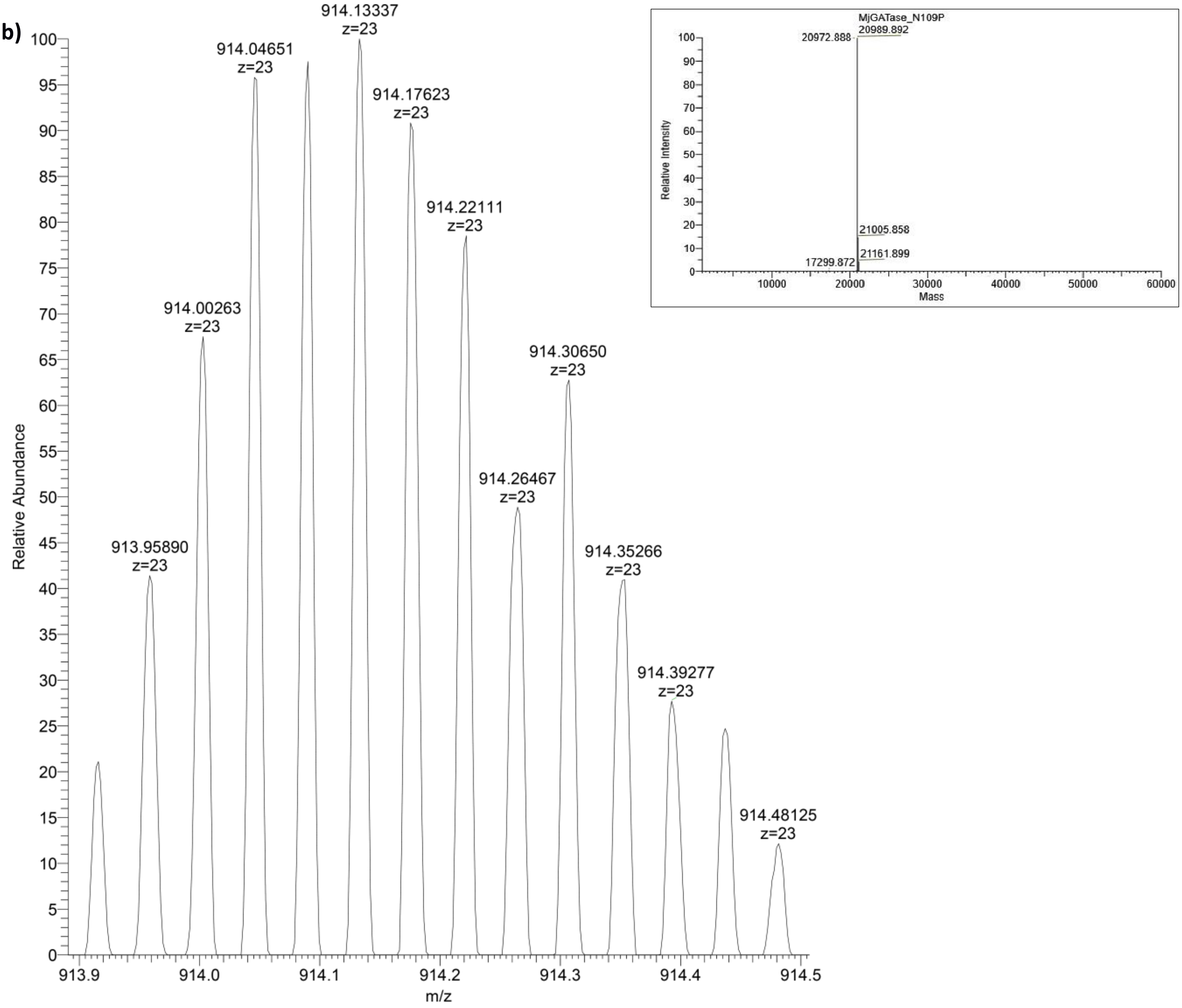

Relative Abundance
m/z
913.95890 z=23
914.00263 z=23
914.04651 z=23
914.13337 z=23
914.17623 z=23
914.22111 z=23
914.26467 z=23
914.30650 z=23
914.35266 z=23
914.39277 z=23
914.48125 z=23
MjGATase_N109P
20972.888
20989.892
21005.858
21161.899
17299.872
Relative Intensity
Mass

**(c)**

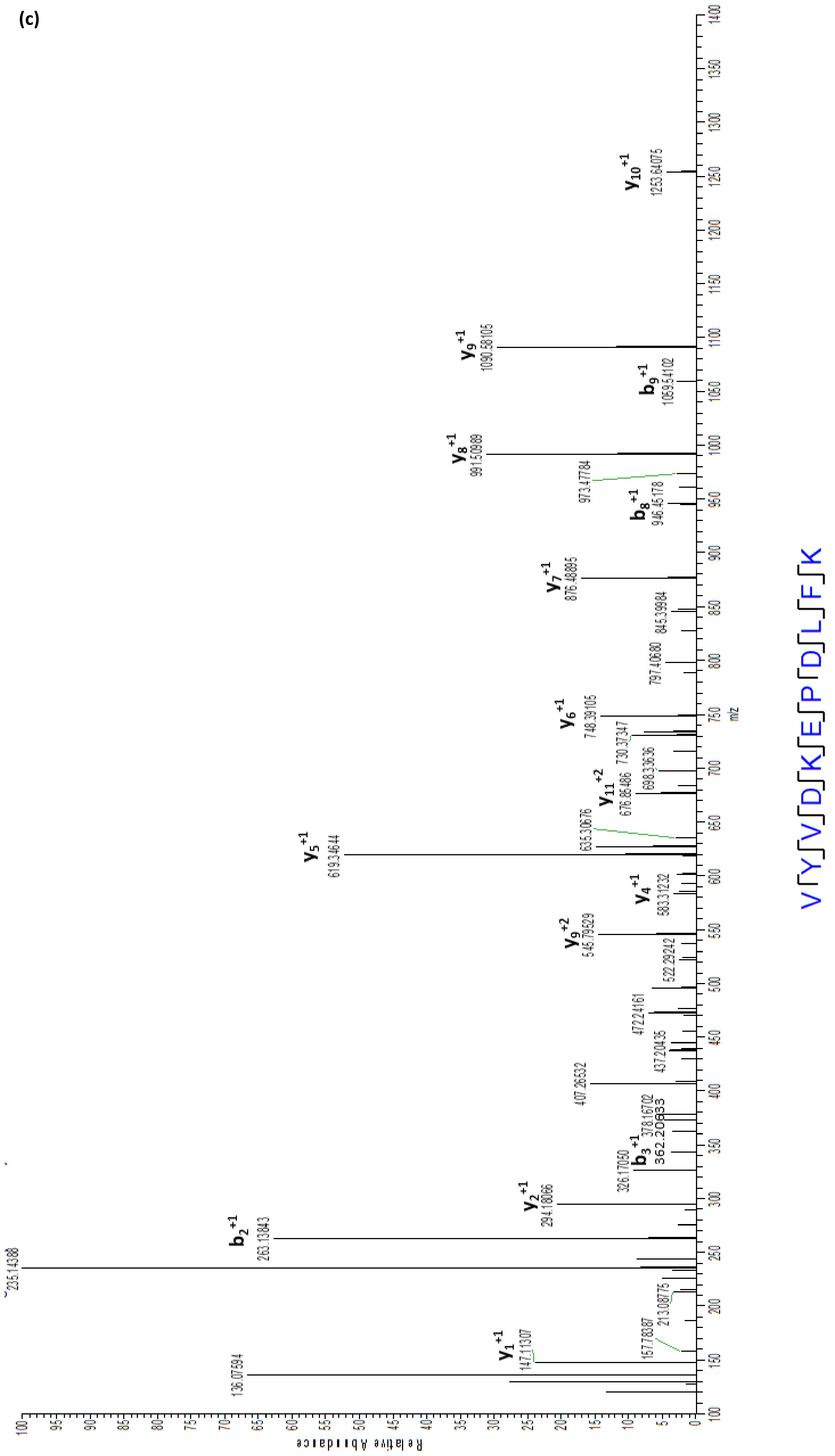
VYVVDKEPDLFK
Relative Abundance
m/z
y1 +1 147.11307
136.07594
157.78387
213.08775
235.14388
b2 +1 263.13843
y2 +1 294.18066
326.17050
b3 +1 362.20633
378.16702
407.26532
437.20435
472.24161
522.29242
y9 +2 545.79529
y4 +1 583.31232
y5 +1 619.34644
635.30676
y11 +2 676.86486
698.33636
730.37347
y6 +1 748.39105
797.40680
845.39984
y7 +1 876.48895
b8 +1 946.45178
973.47784
y8 +1 991.50989
b9 +1 1059.54102
y9 +1 1090.58105
y10 +1 1253.64075

**Supplementary Figure 9:** (a) A portion of DNA sequence chromatogram showing desired mutation from asparagine to proline in MjGATase_SNN109P. (b) ESI-MS spectrum of MjGATase_SNN109P intact protein acquired using Q Exactive HF mass spectrometer. Isotopic mass distribution of the most abundant charged species is shown. Inset shows deconvoluted mass spectrum. The mass of 20989.892 Da corresponds to the expected mass of MjGATase_SNN109P. (c) HCD MS/MS of peptide obtained from in-gel-trypsin digested MjGATase_SNN109P to confirm the presence of prolyl residue in the peptide.

#### Thermal precipitation of proteins

Qualitative evaluation of thermal stability of WT and mutants of MjGATase was done by heating the proteins at five different temperatures. 180 μg of protein (WT, SNN109S and SNN109P MjGATase) was incubated at 75, 80, 85, 90 and 95 °C for 30 minutes and at 100 °C for 15 minutes. The samples were then centrifuged for 10 minutes at room temperature at 24, 000 x g to pellet down precipitated protein fractions. Supernatant corresponding to 90 μg protein was mixed with 4x SDS loading dye of which 20 μg was examined by 12 % SDS-PAGE. At all temperatures, no visible precipitation was seen for WT MjGATase, whereas precipitate was seen at 85 °C for N109S and 80 °C for N109P. At higher temperatures, both mutants continue to show precipitation.

#### Circular Dichroism

CD was performed to compare structure and stability of WT and SNN109S with SNN109P. CD experiments were performed following previously published protocol(1). Briefly, 10 μM of each protein in 5 mM Tris HCl, pH 7.4 and in a cuvette of path length 0.1 cm was used for monitoring thermal unfolding and recording far-ultraviolet (UV) CD (260-200 nm) spectra. Each spectrum was an average of 3 accumulations (scans) and all spectra were corrected for background by subtracting spectrum of the buffer (5 mM Tris-HCl, pH 7.4) from that of the protein. Thermal unfolding experiment was performed at 220 nm by heating the protein from 65 °C to 100 °C with a ramp of 0.5 °C $min^{-1}$. Near-UV (320-260 nm) CD spectra were recorded at 25 °C using 50 μM protein in 5 mM Tris HCl, pH 7.4 and in a cuvette of path length 1 cm. For near-UV measurements, each spectrum was an average of 30 accumulations. The spectrum of the buffer alone was subtracted from all protein spectra.

#### Crystallization and Data collection

Crystallization of WT and mutants were set up under 1:1 mixture of silicon and paraffin oil using the micro-batch method(3). Crystals appeared in less than a week in all the cases. Crystals of MjGATase_WT suitable for X-ray diffraction were obtained at 4 °C in 100 mM sodium acetate pH 4.6 and 35 % PEG 6000 while the crystal of MjGATase_D110G was obtained at 4 °C in 100 mM sodium acetate, pH 5.6 and 10 % PEG 6000 and crystals of MjGATase_SNN109P were obtained in 100 mM sodium acetate, pH 5.6 with 20 % PEG 3350. Prior to diffraction, the crystals of WT and

MjGATase_D110G were soaked in reservoir condition supplemented with 25 % ethylene glycol used as a cryo-protectant for 10 minutes. Crystal of MjGATase_WT diffracted to a resolution of 1.665 Å and a total of 232 frames were collected. The same crystal was then carefully unmounted from the diffractometer and soaked in solution of acivicin for 8 hours and diffracted again under same conditions. It also diffracted at 1.665 Å. Crystal of MjGATase_D110G diffracted at 2.214 Å and 137 frames were collected. Crystal of MjGATase_SNN109P diffracted at 1.89 Å and 180 frames were collected. Details of statistics of data collection and refinement are mentioned in Table 1.

**Structure solution and refinement**

Diffraction images were processed using iMOSFLM(4). The space group of apo and acivicin-bound MjGATase_WT crystals was identified as P32 while that of MjGATase_D110G was $P2_12_12_1$ and MjGATase_SNN109P was P212121.. Structure solutions were obtained using molecular replacement (MR) method using Phaser module of CCP4(5). For WT crystal, structure of GATase from *Pyrococcus horikoshii* (PDB ID: 1WL8) was used as template as it shares 59% identity with MjGATase sequence. Once the WT protein structure was solved, this was used as a MR template for solving structure of acivicin-bound WT, MjGATase_D110G and MjGATase_SNN109P. For refinement of the structures, Refmac module of CCP4 (6, 7) and AutoBuild module of Phenix(8) were used. Refmac placed an asparagine residue at position 109 in the WT structures wherein a clear density for a cyclic residue could be seen. Based on our earlier results(1), we knew that N109 is cyclized to succinimide (SNN) in native MjGATase. Hence, N109 was replaced by SNN, the coordinates of which were available at J ligand library, during phasing and then several rounds of refinement were carried out using Refmac. After refinement, SNN fitted correctly within the electron density and was seen to form appropriate covalent bonds with E108 and D110. In case of liganded structure, co-ordinates for acivicin (5CS) were also procured from J ligand and included in the structure co-ordinates file and multiple rounds of refinement were carried out then on. In case of MjGATase_D110G, the electron density corresponding to E108-D110 tripeptide segment was found to be missing, while the rest of the protein showed a good electron density map (Supplementary Figure 1c). The structure of MjGATase_D110G is reported with region between E108-D110 unmapped. In case of MjGATase_SNN109P, the electron density corresponding to residues 93-95 was found to be missing in chain A out of the 4 chains in the asymmetric unit.

**B) Computational Methods**

**Regular Molecular Dynamics (MD) simulations**

MD simulations were performed using GROMACS 5.1.4(9, 10) with GROMOS54a7 united-atom force field(11) parameters for the protein and SPCE model(12) for water molecules. See subsection i for the details about the force field parameterization for succinimidyl residue. The enzyme (MjGATase_WT MjGATase_SNN109S or MjGATase_SNN109P) was placed at the centre of a cubic box with its edges away from the protein surface by at least 12 Å. The enzyme was then solvated using PACKMOL(13), followed by addition of ions to neutralize the system. A steepest descent energy minimization was performed keeping position restraints on heavy atoms of the enzyme with a force constant of 4184 kJ/mol/rad$^2$. This minimized structure was then equilibrated in multiple steps. Firstly, a 300 ps NVT (at 300 K) followed by a 500 ps NPT equilibration run (at 300 K and 1 bar) were carried out with position restraints on the heavy atoms of the enzyme with a force constant of 4184 kJ/mol/rad$^2$. The NPT run was extended further for a duration of 300 ns by removing all position restraints. The Bussi-Donadio-Parrinello velocity rescaling thermostat(14) with coupling constant 0.5 ps at 27 °C was used in the NVT runs and the same thermostat and Parrinello-Rahman barostat(15, 16) with coupling constant of 1.0 ps for both, at 300 K and 1 bar respectively were used in the NPT runs. An integration time step of 2 fs was used along with LINCS constraints(17) on all bonds. Particle mesh Ewald (PME) method(18) with cut-off distance of 14 Å was used to treat the long-range electrostatic interactions.

We have performed normal MD simulations for WT enzyme and its SNN109S and SNN109P mutants at 27, 70 and 87 °C for 300 ns each. But these results do not show any significant difference in either the structure or dynamics between WT and SNN109S mutant (see subsection i), due to inadequate sampling of conformations. Following the above protocol, normal MD simulation was also performed at 27 °C for the MjGATase_D110G mutant and PhGATase (an enzyme homologous to MjGATase, PDB ID: 1WL8).

**Replica exchange with solute scaling (REST2) simulations**

Within MD simulation time scales, MjGATase (WT) and succinimide lacking mutant MjGATase_SNN109S and MjGATase_SNN109P do not show any difference in stability due to inadequate sampling of conformations. Therefore, to enhance conformational sampling, we performed replica exchange with solute (enzyme) scaling simulations (REST2)(19, 20). REST2 is an improved type of Hamiltonian replica exchange method. Instead of tempering the whole system (like normal Hamiltonian replica exchange simulation(21)), this REST2 method specifically scales the potential energy surface of the region of interest (solute) resulting in a significant decrease in the number of replicas; thus making it computationally effective and enabling the study of systems without an a priori knowledge of major conformational change(s), i. e., the unfolding of small proteins with high energy barrier(22–25), high energy driven conformational changes of large proteins(26, 27) and ligand-binding to proteins(25, 27–32). Herein, REST2 simulations were performed using GROMACS 5.1.4 patched with PLUMED 2.3.0(33, 34), separately for the WT and its SNN109S and SNN109P mutants (starting from the last frame of the corresponding normal MD simulation) with 28 replicas for each system in the effective temperature range 47-327 °C (320-600 K) for the tempered region (protein) while the solvent was still in reference temperature of 47 °C (320 K) for all replicas. The replicas were allowed to exchange every 10 ps. Rest of the simulation protocol was similar to what was described for the standard MD simulations. The average exchange probability between the successive replicas was found to be 35% (see subsection i for details).

**Regular MD after REST2**

After 250 ns long REST2 simulation with 28 replicas, the last frame of the fourth and sixth replicas (with REST2 "effective" temperatures of 70 and 87 °C, respectively) were chosen and warmed up gradually (through simulated annealing) to 70 and 87 °C, respectively. Normal MD (NPT ensemble) was then performed for 300 ns for these systems at these temperatures. The analysis presented in this manuscript in Figure 3 (f and g), Figure 4 and Figure 5 (b and c) from the main text and Supplementary Figures 5b and 7 (except from the results related to 27 °C) are from these 200-300 ns portion of the normal MD trajectories, post the REST2 run.

**Well-tempered metadynamics (WT-MTD) simulations**

The key results obtained from the REST2 simulations are validated further by well-tempered metadynamics, a popular biased enhanced sampling technique(35). The free energy profile for WT MjGATase and its SNN109S mutant with respect to the radius of gyration (collective variable or CV) of the residues from the succinimide containing loop (residues 108-115) was reconstructed from 2200 ns long well-tempered metadynamics simulations (WT_MTD) at 27 °C (300 K). The WT-MTD simulations were performed with a well-tempered factor (bias factor) of 8. An initial deposition rate of 2.5 kJ/mol/ps for the biasing Gaussian potential of width 0.016 Å (for WT) and 0.027 Å (for SNN109S mutant) was used. The width of the Gaussian bias was determined based on the fluctuation of CV with simulation time calculated from the normal MD simulation of the corresponding system. The integration timestep for WT-MTD simulation was 1 fs. All other parameters are same as in the normal MD simulation. The radius of gyration $R_g$ (collective variable) was calculated as follows:

Radius of gyration, $R_g = \left(\frac{\sum_i m_i (r_i - r_{COM})^2}{\sum_i m_i}\right)^{\frac{1}{2}}$

where, $r_{COM} = \frac{\sum_i m_i r_i}{\sum_i m_i}$

Here $r_i$ and $m_i$ are the position and mass of the $i$-th atom; and $r_{COM}$ is the centre of mass of all atoms present in residues 108-115.

Flow chart representing the entire MD simulation protocol followed in the present study in shown in Supplementary Figure 16. See the validation of these computational methods in the subsection i.

**Natural Bond Orbital (NBO) analysis**

The initial structure of peptide segment 108-112 was extracted from the crystal structure. All other residues except SNN109 were truncated to alanine, and the N and C termini were acetylated and amidated, respectively. Density functional theory (DFT) calculations was performed at the B3LYP/6-311++G(2d,p) level with NBO analysis(36) using second-order perturbation theory to estimate the contributions of the lone pairs on the backbone carbonyl oxygen of Glu 108 to the n→π* interaction and n→σ* interaction (hydrogen bond). This calculation was carried out with NBO 3.1(37) as implemented in Gaussian 09(38). The threshold for printing the second-order stabilization energies was set to 0.04 kJ/mol. The figures representing orbital overlap were generated with Chemcraft(39) .

**Analysis method**

Trajectories were visualized using VMD(40). PyMOL(41) and VMD were used to prepare the graphics. Results were obtained from the analysis of the trajectories of normal MD at 27 °C, 70 °C (started from last frame of fourth replica in REST2) and 87 °C (started from last frame of sixth replica in REST2) within 200-300 ns simulation time window. For WT-MTD simulations, the entire 2200 ns trajectory was used to reconstruct the free energy profile. All the figures presented as snapshots from MD simulation both in the main text as well as in the SI were taken from the conformation of the central member of the most predominant cluster obtained from cluster analysis carried out using GROMOS algorithm(42) (Supplementary Table 2). Modules in-built in GROMACS and PLUMED as well as home-grown scripts were used for analyses.

**Supplementary Table 2:** Detailed statistics of cluster analysis.

| System | Choice of frames | Number of frames | RMSD cut-off (Å) | % contribution of predominant cluster |
|---|---|---|---|---|
| Normal MD, WT, 27 °C | 200-300 ns | 10,000 | 1.7 | 87.10% |
| Normal MD post-REST2, WT, 70 °C | 200-300 ns | 10,000 | 1.9 | 74.80% |
| Normal MD post-REST2, WT, 87 °C | 200-300 ns | 10,000 | 2.3 | 90.10% |
| Normal MD, SNN109S, 27 °C | 200-300 ns | 10,000 | 1.8 | 72.60% |
| Normal MD post-REST2, SNN109S, 70 °C | 200-300 ns | 10,000 | 2.1 | 75.00% |
| Normal MD post-REST2, SNN109S, 87 °C | 200-300 ns | 10,000 | 3.1 | 75.00% |
| Normal MD, SNN109P, 27 °C | 200-300 ns | 10,000 | 1.5 | 84.40% |
| Normal MD post-REST2, SNN109P, 70 °C | 200-300 ns | 10,000 | 1.7 | 61.80% |
| Normal MD post-REST2, SNN109P, 87 °C | 200-300 ns | 10,000 | 2.2 | 87.70% |
| WT-MTD, WT, 27 °C, free energy minimum | Rg: 5.36-5.56 Å | 27,671 | 2.2 | 83.70% |
| WT-MTD, SNN109S, 27 °C, lower free energy minimum | Rg: 5.52-5.72 Å | 18,550 | 2.5 | 76.10% |
| WT-MTD, SNN109S, 27 °C, higher free energy minimum | Rg: 6.90-7.10 Å | 6,665 | 2.5 | 67.40% |

### i) Validation of Computational Methods

#### a) Force Field parameterization for succinimide residue

The structure of the succinimide residue (SNN) with acetate (Ac) and NMe caps (Ac-SNN-NMe Supplementary Figure 10) was taken from the crystal structure of MjGATase. The force field parameters for SNN was extracted from gromos54a7 (under GROMOS96) united atom force field (FF) by following the parameterization rules prescribed by Petrov *et al*(43) who had developed force field parameters (GROMOS) of more than 250 different types of enzymatic and non-enzymatic post-translational modifications (PTMs) and validated these against the corresponding experimental hydration free energies. In this regard, we also took help of an intuitive structural analogy of succinimide to pentagonal ring containing residues (proline and pyroglutamic acid) and N5-methylglutamine (Supplementary Figure 10) whose force field parameters were already reported within an extended list of gromos54a7 parameters tested and provided by Vienna-PTM web server(44) (http://vienna-ptm.univie.ac.at/). Herein, we have benchmarked these FF parameters for SNN against the quantum optimized structure. Towards this purpose, the Ac-SNN-NMe structure was optimized in gas phase using Gaussian09 package(38) at B3LYP/6-31+g(d,p) level of theory. The initial configuration of Ac-SNN-NMe for this quantum optimization was obtained from the crystal structure of MjGATase (the residues 108-110 containing SNN109 was taken from the crystal structure followed by deletion of atoms from E108 and D110 to convert them to Ac and NMe caps (to SNN), respectively). The classical MD simulation with gromos54a7 FF was performed for Ac-SNN-NMe (in both gas phase as well as in water) and also the whole MjGATase protein in water at 27 °C (and 87 °C) and 1 bar using the same protocols used in the simulations of WT and SNN109S and SNN109P mutants as described in the main text.

From Supplementary Table 3 and Figure 11 (left), it is seen that the potential energy profile of the C-$C_\alpha$-$C_\beta$-$C_\gamma$ dihedral angle of SNN has two minima, one at 12° and another at -14°. This is quite expected as a pentagonal ring can have two different conformations other than planar (planar form is generally not stable), viz. puckered-up and puckered-down. This is also supported by the distribution of similar dihedral present in proline (which also contains a pentagonal ring) as shown in Supplementary Figure 11 (right) for P30 and P164 in MjGATase. Supplementary Figure 12 shows that all the optimized structures of Ac-SNN-NMe both from quantum (in gas phase) and classical (both in gas phase and in water) level of optimization are close to the configuration of Ac-SNN-NMe obtained from the crystal structure of MjGATase, thus validating the force field parameters.

The relaxed potential energy profile scan with respect to the C-$C_\alpha$-$C_\beta$-$C_\gamma$ dihedral of SNN within the puckered-up and puckered-down region of SNN pentagonal ring (-30 to 30°) shows that these profiles calculated at both classical and quantum levels are comparable (Supplementary Figure 13).

Finally, the overlay of residues 108-110 (containing residue SNN109) between MjGATase crystal structure and the structure obtained after 300ns of MD simulation of MjGATase (at 27 °C and 1 bar) using the developed force field parameters for SNN reveals that these parameters can hold the MjGATase structure close to its crystal structure (Supplementary Figure 14).

All these observations support the use of the developed force field parameters for SNN residue in MD simulation of MjGATase. These force field parameters for both capped-SNN (Ac-SNN-NMe) and only SNN residues are documented in subsection ii.

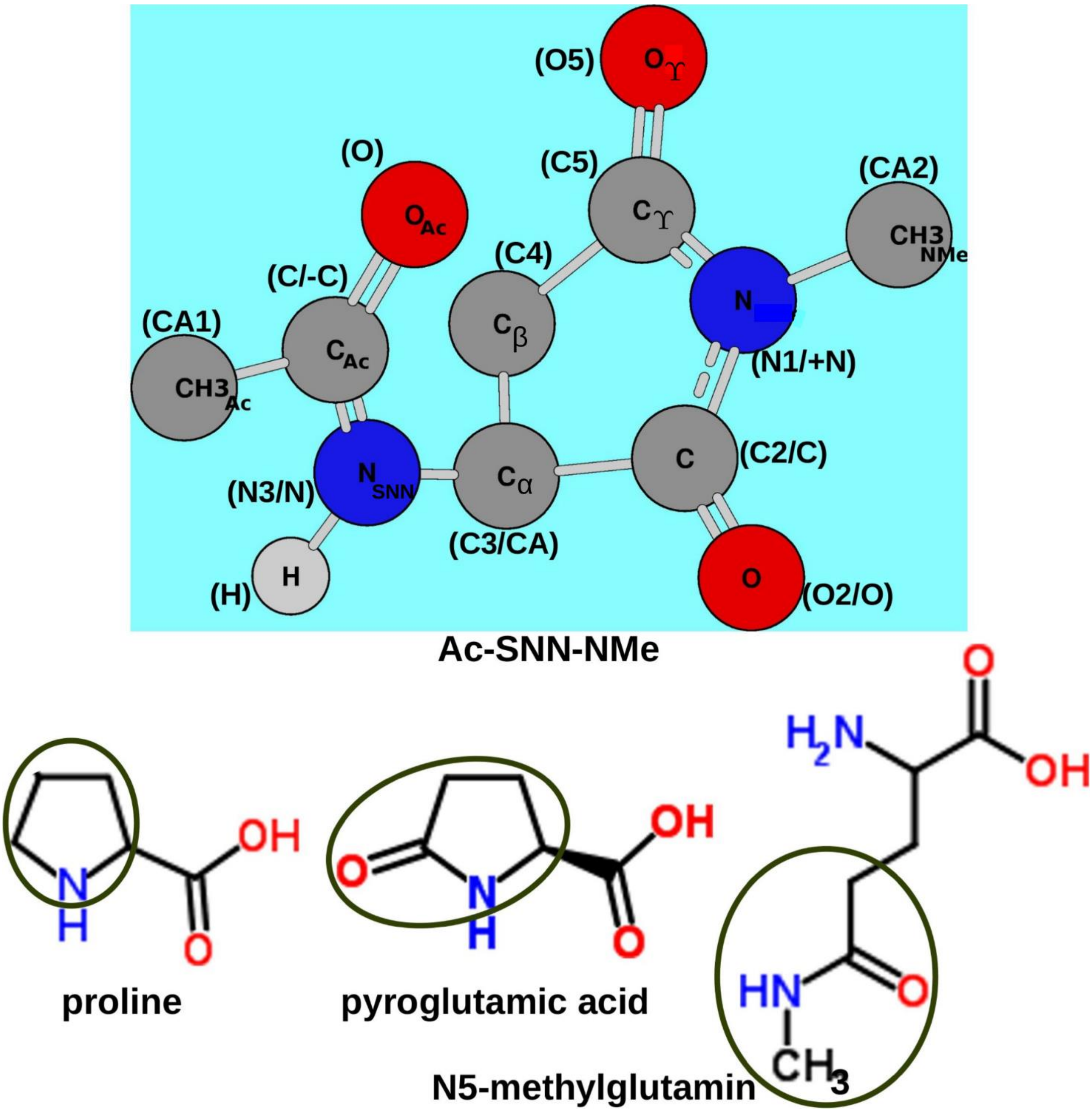


**Supplementary Figure 10: Above:** Structure of Ac-SNN-NMe extracted from MjGATase crystal structure (The residues 108-110 are extracted from MjGATase structure containing SNN. Except from $C_\alpha$, C and O, rest of the atoms of residues 108 are deleted and $C_\alpha$ is capped with three hydrogen atoms. Similarly, except from N and $C_\alpha$, rest of the atoms of residues 110 are deleted and $C_\alpha$ is capped with three hydrogen atoms. Thus, we generated the coordinates for the atoms in Ac-SNN-NMe.). This structure of Ac-SNN-NMe is used to benchmark the force field parameters of SNN with respect to quantum calculations. Name of each atom is written within circle representing the corresponding atom. Name of each atom according to force field are written within bracket. The atoms for those the name according to force field is different between capped and uncapped SNN (i. e., Ac-SNN-NMe and only SNN) are represented with the format: ($name_{capped}$/$name_{uncapped}$). The force field parameters for both capped-SNN (Ac-SNN-NMe) and only SNN residues are documented later. **Below:** The molecules whose force field parameters were already reported within an extended list of gromos54a7 parameters tested and provided by Vienna-PTM web server and used to extract the force field parameters for SNN by structural analogy (similar portion with respect to SNN is shown within ellipses).

**Supplementary Table 3:** Dihedral angle C-$C_\alpha$-$C_\beta$-$C_\gamma$ of SNN at different levels of computation (quantum and classical) starting from various initial structures.

| Initial structure | C-$C_\alpha$-$C_\beta$-$C_\gamma$ dihedral angle of SNN (degree) | | |
|---|---|---|---|
| | DFT B3LYP/6-31+g(d,p) (in gas phase) | Classical MD | |
| | | In gas phase | In water |
| Ac-SNN-NMe (crystal str.) | +12.0 | -13.2 | +12.0 |
| after 40 ns of NPT | +18.1 (residues 108-110) | -- | -14.0 (whole protein) |
| Ac-SNN-NMe (B3LYP optimized) | -- | +11.4 (gas phase) | -- |

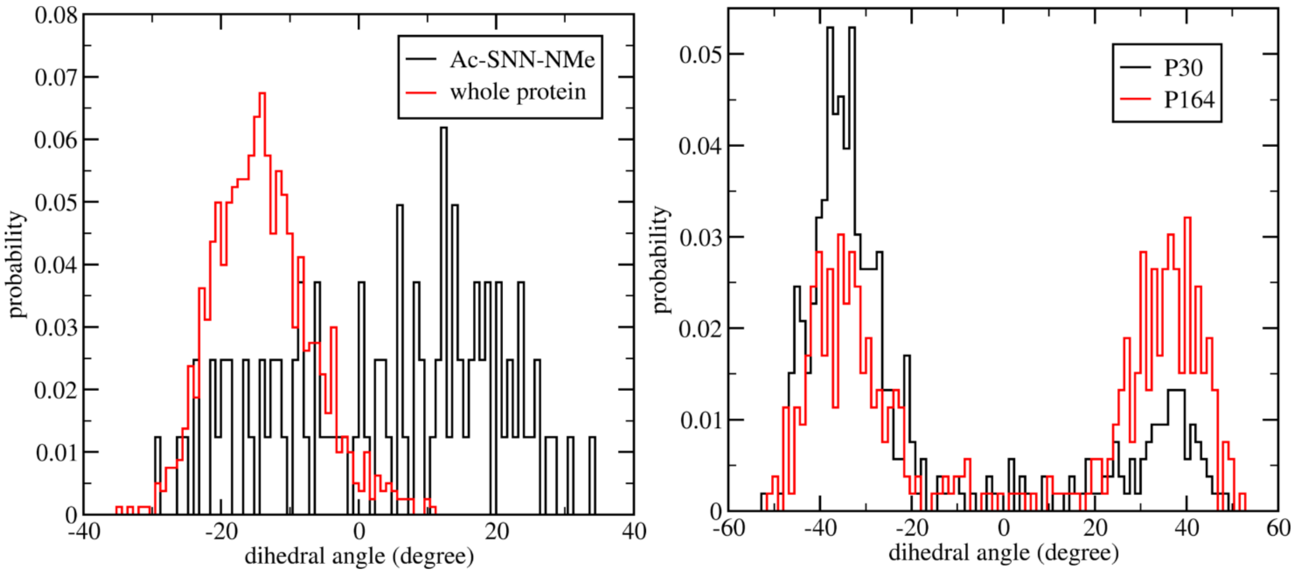


**Supplementary Figure 11: Left:** Distribution of C-$C_\alpha$-$C_\beta$-$C_\gamma$ dihedral angle of pentagonal ring of SNN from the NPT simulation of Ac-SNN-NMe (at 27 °C) and whole MjGATase (at 27 °C and 87 °C) in water. The value of the corresponding dihedral angle in MjGATase crystal structure is found to be +8.30$^0$. **Right:** Distribution of C-$C_\alpha$-$C_\beta$-$C_\gamma$ dihedral angle of pentagonal ring of P30 and P164 from the NPT simulation of MjGATase. The value of the corresponding dihedral angle in crystal structure is found to be -35.50$^0$ for both P30 and P164.

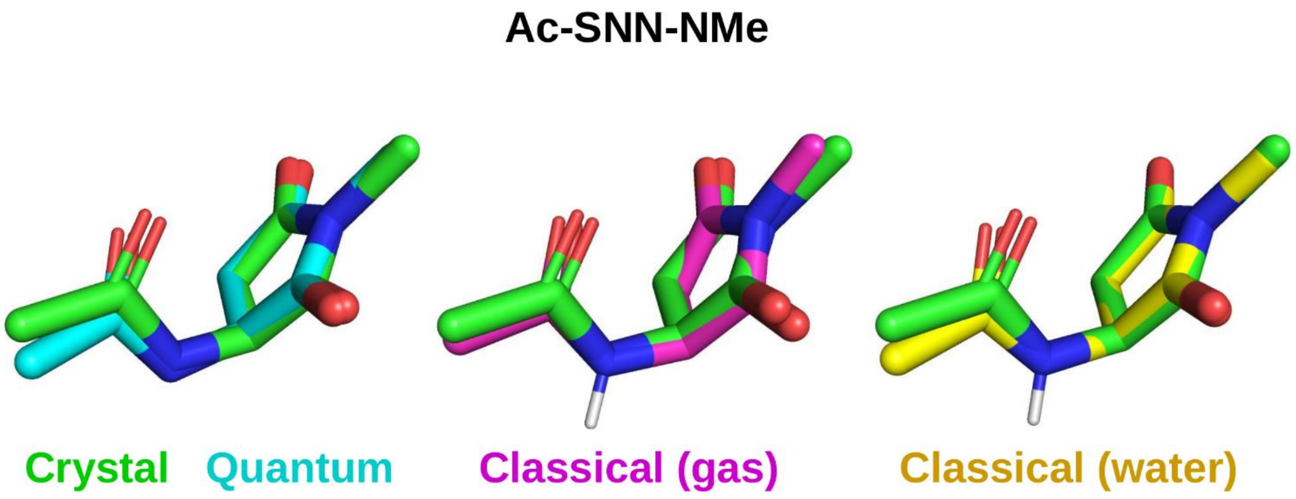


**Supplementary Figure 12:** Overlay of Ac-SNN-NMe both from quantum (in gas phase) and classical (both in gas phase and in water) optimized structure with the configuration of Ac-SNN-NMe obtained from the crystal structure of MjGATase.

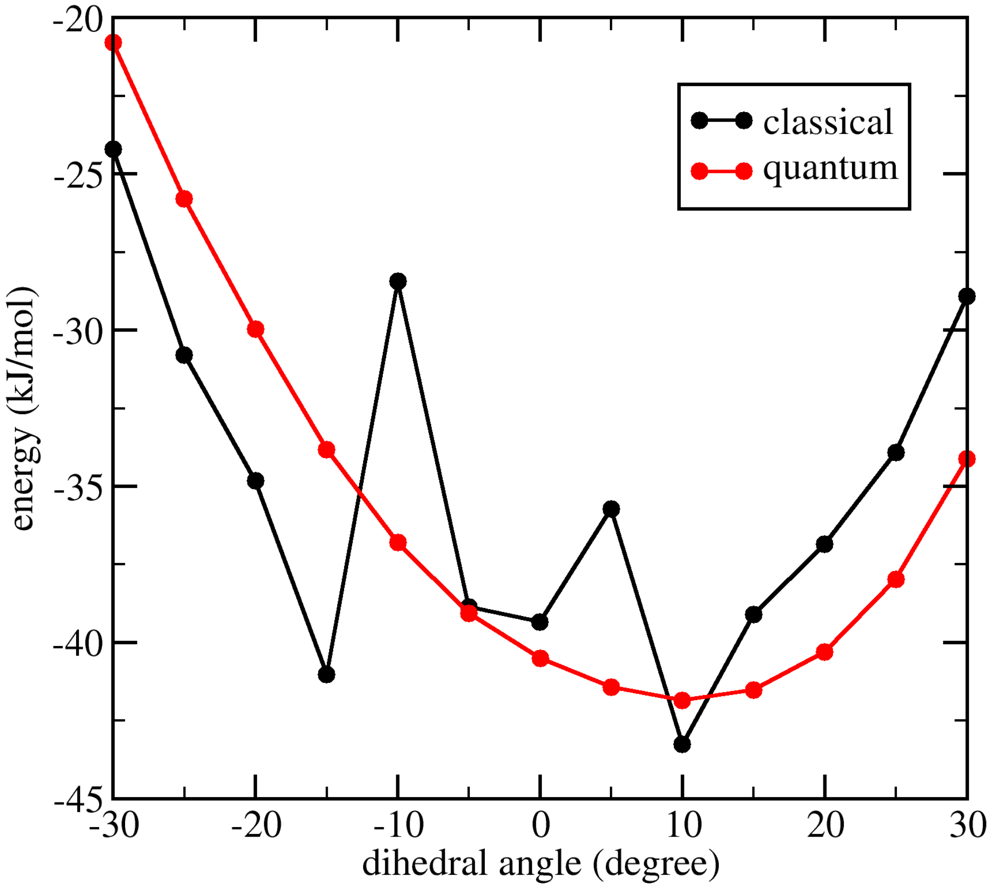


**Supplementary Figure 13:** Relaxed potential energy profile scan for the C-$C_\alpha$-$C_\beta$-$C_\gamma$ dihedral of SNN.

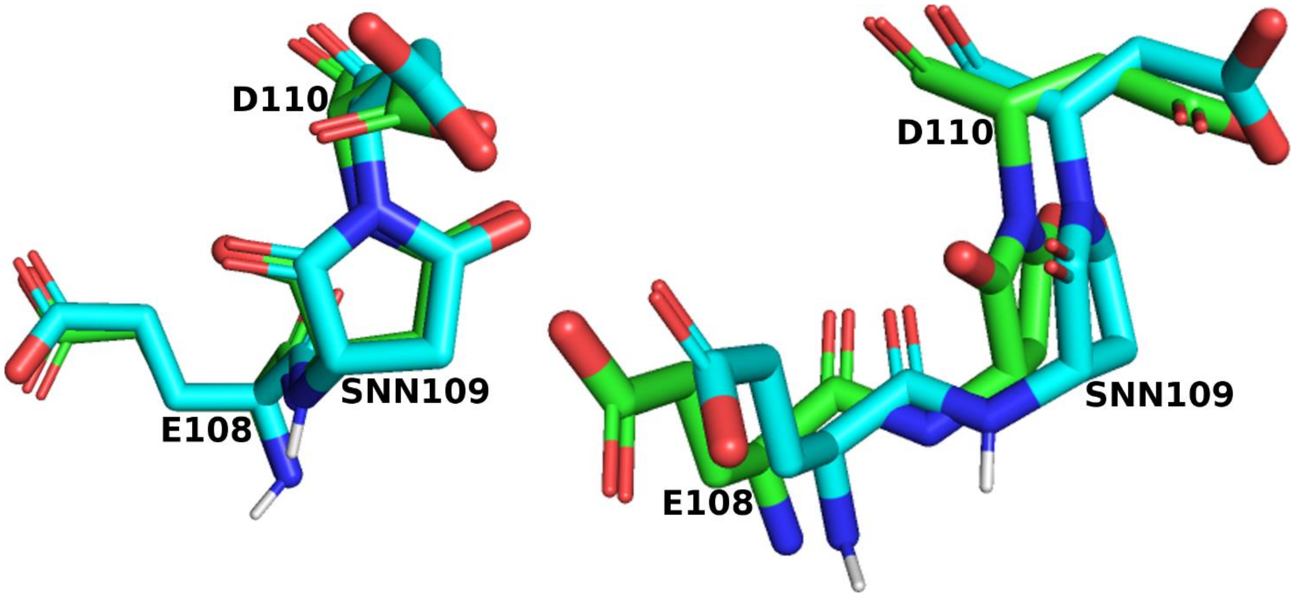


**Supplementary Figure 14: Left:** Overlay of residues 108-110 (containing residue SNN109) between MjGATase crystal structure (green) and the structure obtained after 300ns of MD simulation of MjGATase at 27 °C and 1 bar (cyan). **Right:** Rotated view of the left panel.

#### b) Validation of REST2 runs

According to the REST2 method, the fluctuation of Epp + $(1/2)(\beta_0/\beta_m)^{1/2}E_{pw}$ (where, Epp and $E_{pw}$ are intra-protein and protein-solvent interaction energies, respectively; and $\beta_0 = 1/(k_BT_m)$, where, $k_B$ is Boltzmann constant and $T_m$ is the effective temperature of the m-th replica) can be thought to determine the acceptance ratios for exchanges between the m-th replica and its neighboring replicas(45). A considerable degree of overlap between the distributions of the energy term Epp + $(1/2)(\beta_0/\beta_m)^{1/2}E_{pw}$ between neighboring replicas for a 250 ns (per replica) long REST2 simulation run for MjGATase_WT, MjGATase_SNN109S and MjGATase_SNN109P mutants (Supplementary Figure 15) leads to 35% average exchange probability among the neighboring replicas for each of these systems. This validates the protocol used for our REST2 simulations.

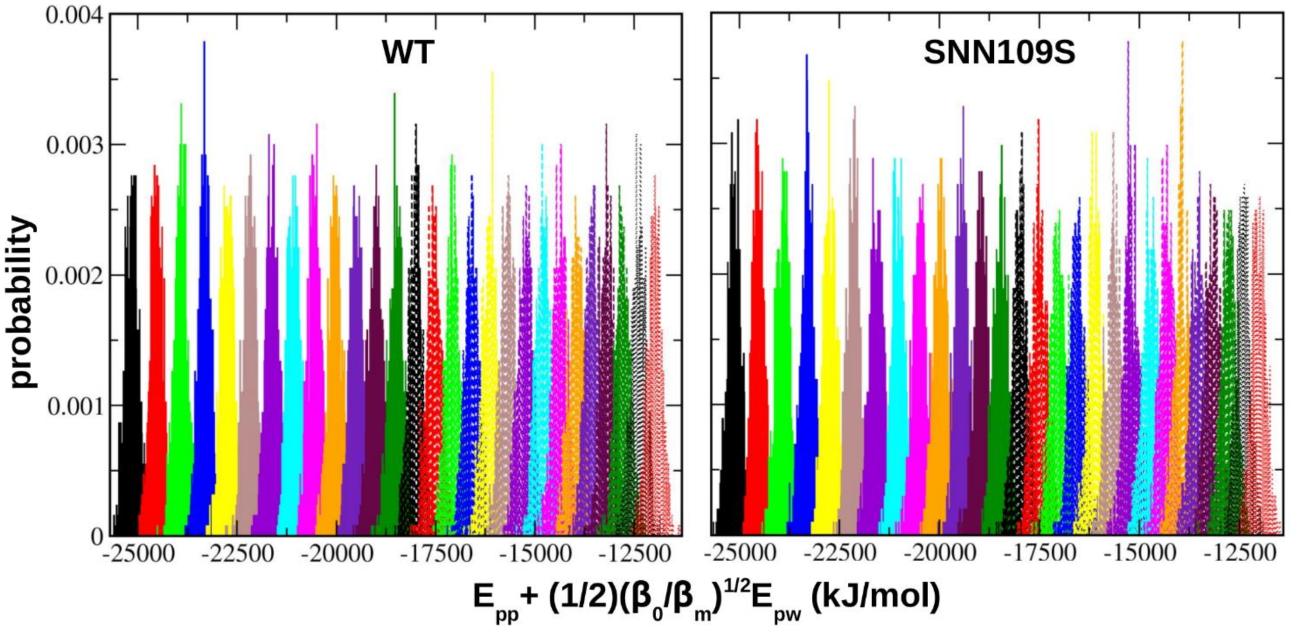


**Supplementary Figure 15:** Distribution of the energy term Epp + $(1/2)(\beta_0/\beta_m)^{1/2}E_{pw}$ between neighboring replicas for a 250 ns (per replica) long REST2 simulation run for WT (left) and its SNN109S (right) and SNN109P mutants (bottom). These distributions start from the first replica (solid black) from the left and end at last (28-th) replica (dotted red) at the right.

#### c) Necessity of REST2: Enhanced sampling over normal MD

Now, we inspect whether REST2 simulations help in enhancing the conformational sampling of MjGATase over normal MD or not. At this point, for the ease of understanding, the entire simulation protocol has been explained with the help of a flow chart (Supplementary Figure 16).

A 250 ns (per replica) long REST2 simulation run with 35% average exchange probability results in the spread of around 70% of the replicas throughout the overall REST2 effective temperature space (47-327 °C) for both WT (Supplementary Figure 17) and SNN109S mutant (Supplementary Figure

18). This ensures a considerable enhancement in the conformational sampling of MjGATase through REST2 simulation over normal MD simulation (However, in a REST2 simulation, except the lowest replica, all other replicas are not present in any specific thermodynamic ensemble for MjGATase.). Thus, a protocol made of REST2 followed by normal MD at the respective temperature was adopted (Supplementary Figure 16). This procedure successfully demonstrates the conformational difference of MjGATase between its WT and mutated forms as revealed by the conformation of the extended β-hairpin (Supplementary Figure 19), RMSF (Supplementary Figure 20) and fraction of native contacts (Supplementary Figure 21) of MjGATase in its WT and mutated forms. While the overlay of extended β-hairpin regions between WT and SNN109S and SNN109P mutants (Figure 5c in the main text) and the RMSF plot for WT and SNN109S and SNN109P mutants (Supplementary Figure 7) obtained from Run B simulations (Supplementary Figure 16) exhibited significant difference, those obtained from Run A (Supplementary Figure 16) simulations do not (Supplementary Figures 19 and 20). This observation brings forth the vital role of REST2 based sampling.

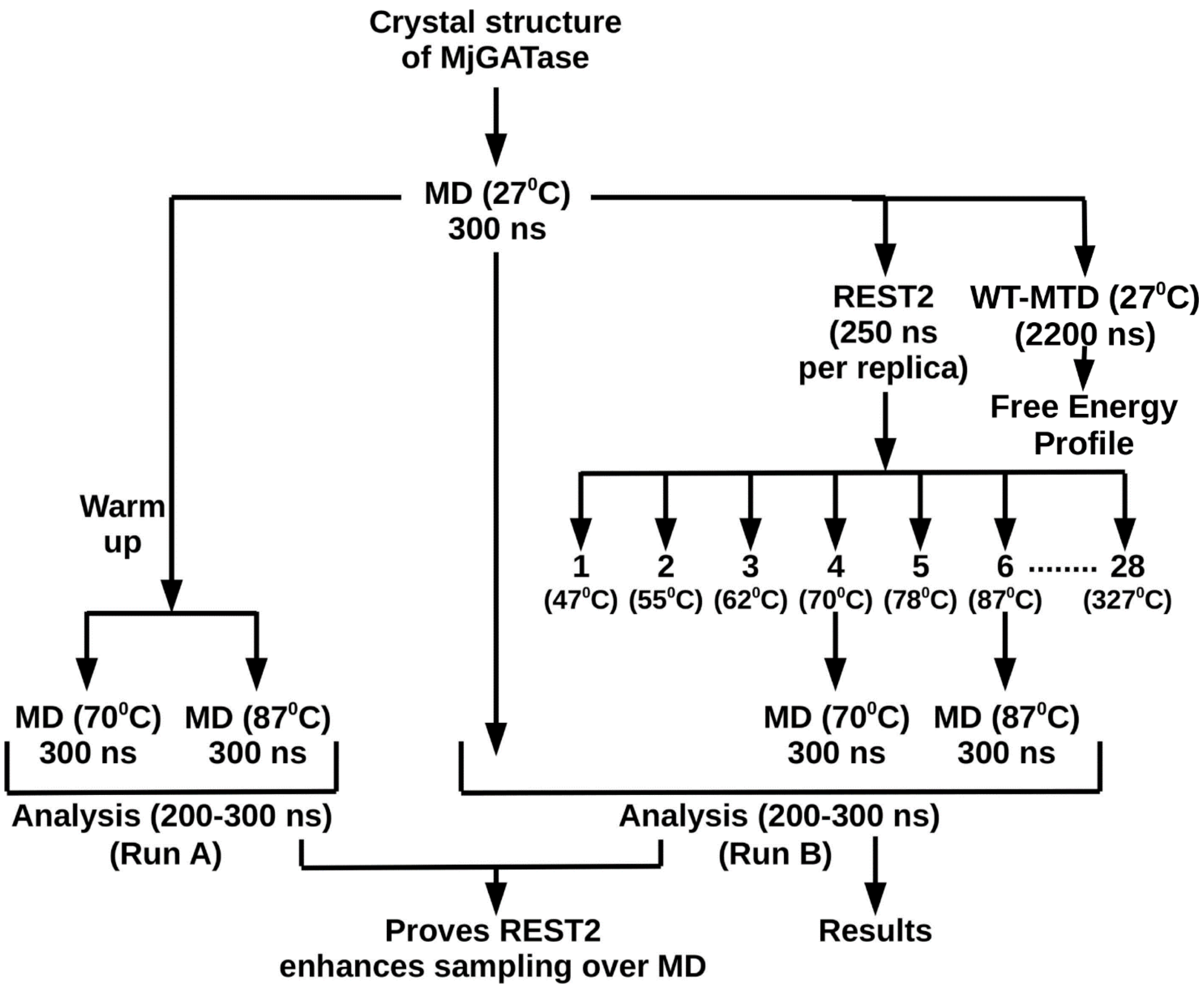

**Supplementary Figure 16:** Flow chart representing the entire MD simulation protocol followed in the present study of WT enzyme. Same protocol was followed for the SNN109S mutant as well. Regular MD and REST2 simulations were performed for SNN109P mutant, following the same protocol. For MjGATase_D110G and PhGATase_WT, 300 ns long normal MD simulation have been performed. Overall, more than 31 microseconds of atomistic MD simulations have been carried out for this project.

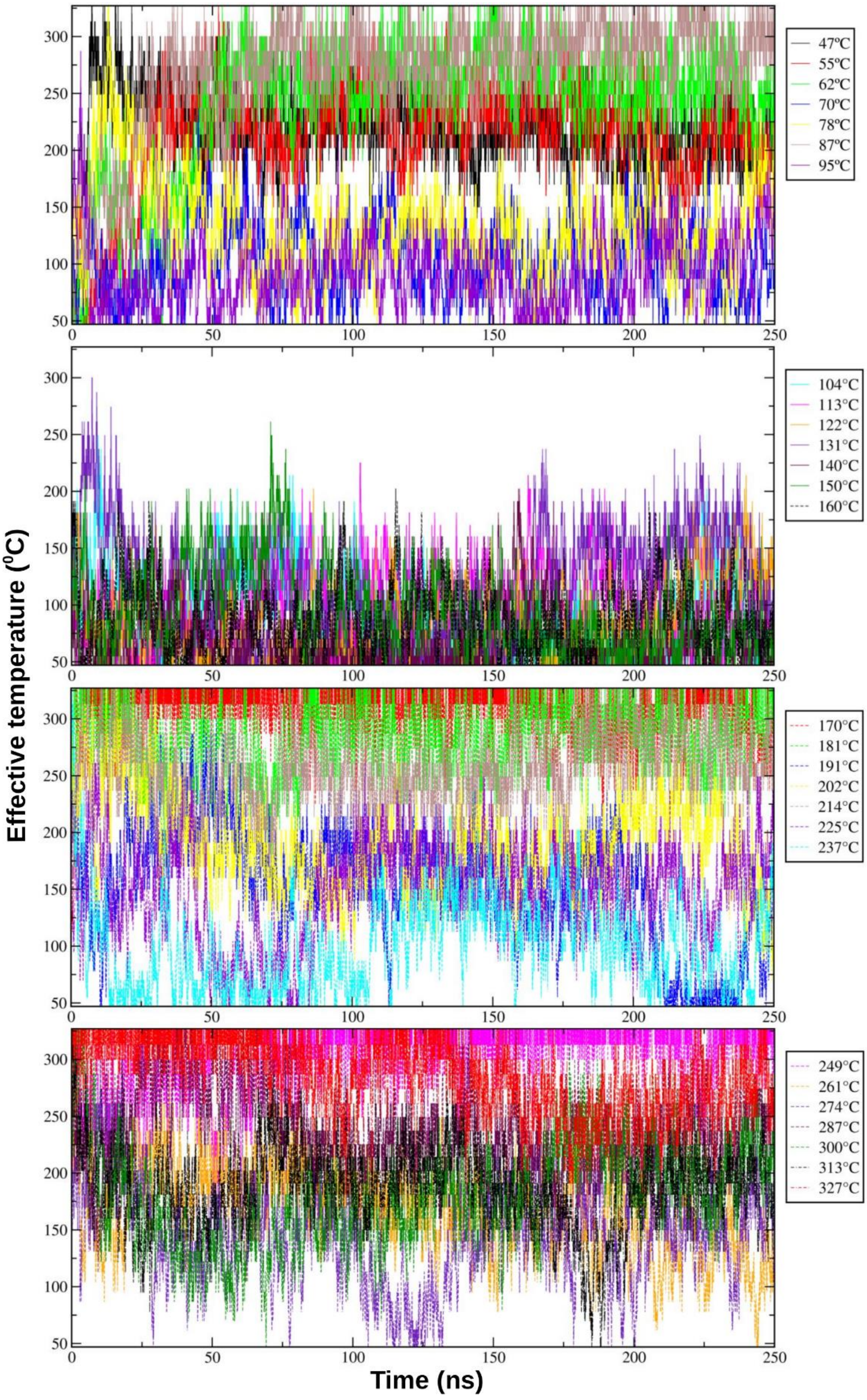

**Supplementary Figure 17:** The spread of each replica (represented as their corresponding effective temperatures in the legend boxes) over the REST2 effective temperature space (47-327 °C) in a 250 ns (per replica) long REST2 simulation of the WT enzyme. Each graph contains seven successive replicas.

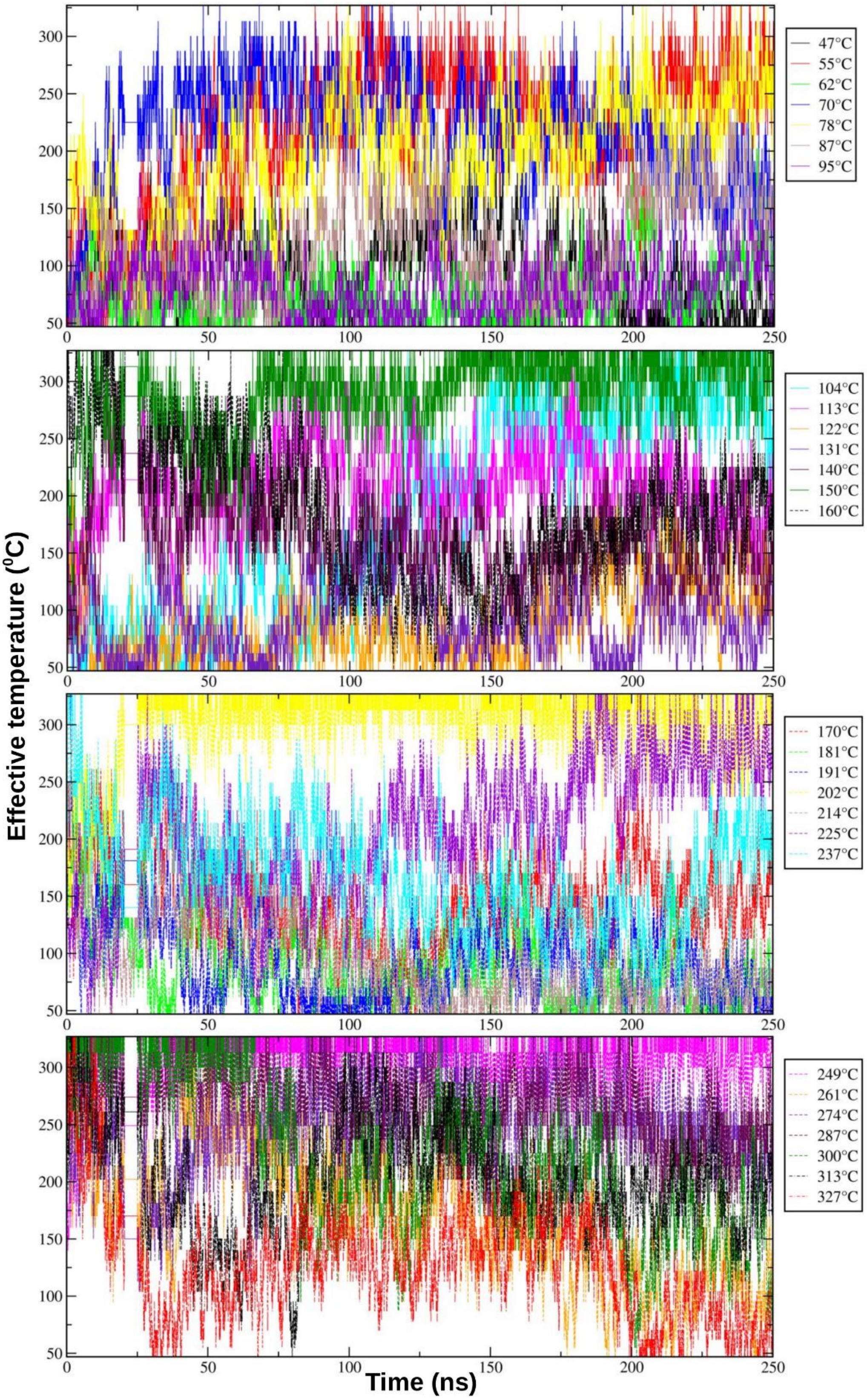


**Supplementary Figure 18:** The spread of each replica (represented as their corresponding effective temperatures in the legend boxes) over the REST2 effective temperature space (47-327 °C) in a 250 ns (per replica) long REST2 simulation of the SNN109S mutant. Each graph contains seven successive replicas.

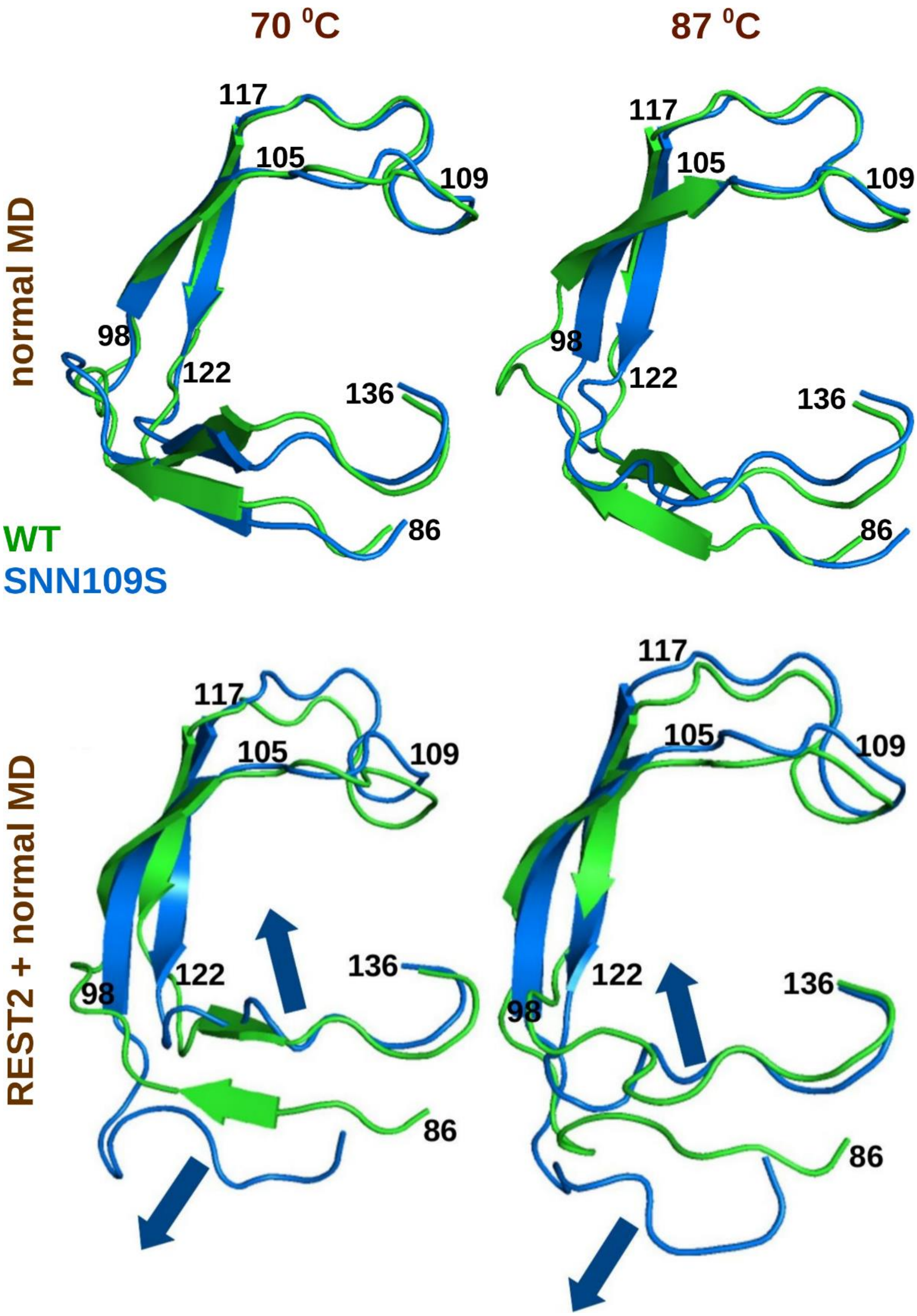


**Supplementary Figure 19:** Comparison of the overlay of extended β-hairpin region (residues 86-136) of MjGATase in its WT (green) and SNN109S mutant (blue) forms between only normal MD (Run A, see Supplementary Figure 16) and combined REST2 and normal MD approach (Run B). The conformation of the central member of the most predominant cluster resulted from cluster analysis of 200-300 ns trajectory of the corresponding simulation has been presented here.

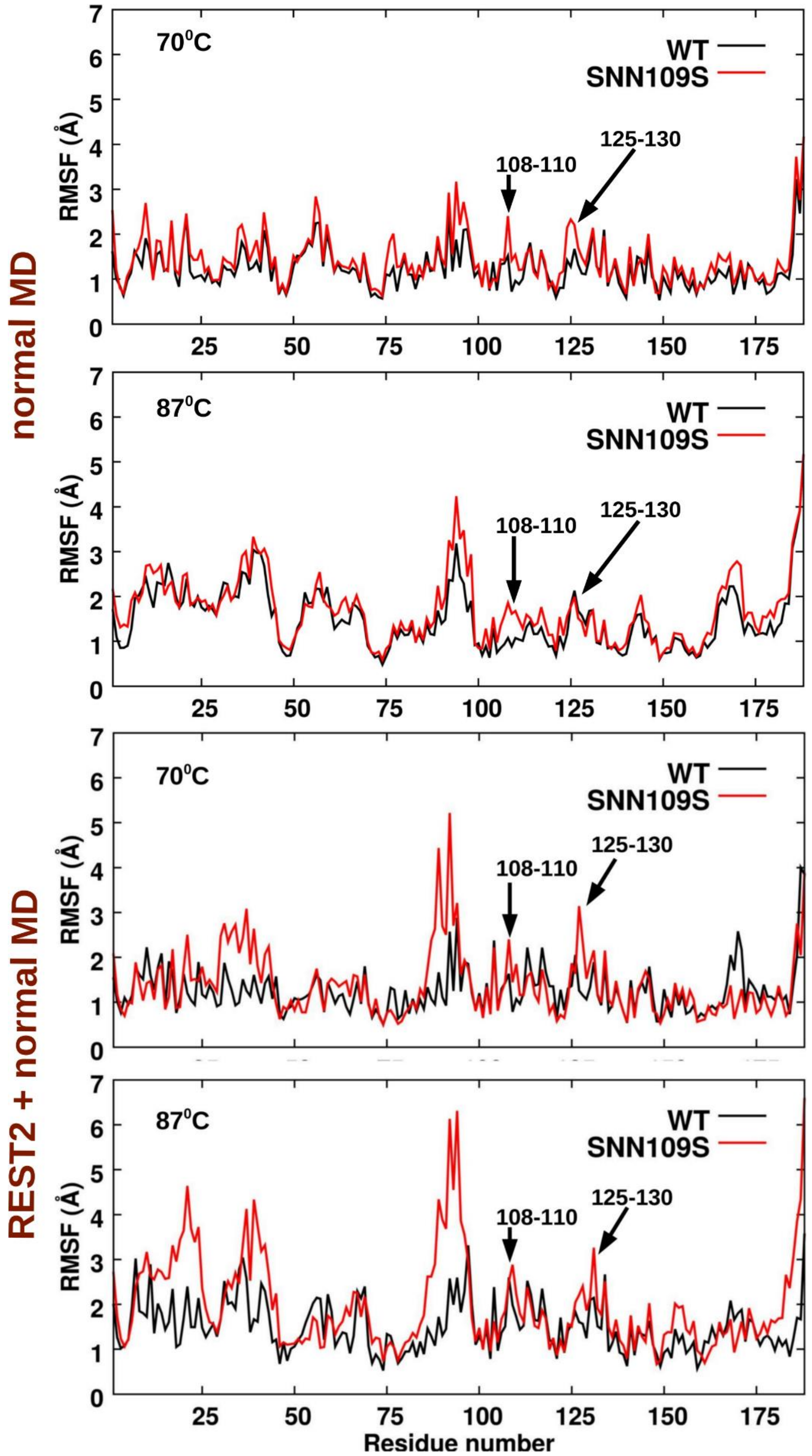


**Supplementary Figure 20:** Comparison of the root mean square fluctuations (RMSF) of residues of MjGATase in its WT and SNN109S mutant forms between only normal MD (Run A, see Supplementary Figure 16) and combined REST2 and normal MD approach (Run B).

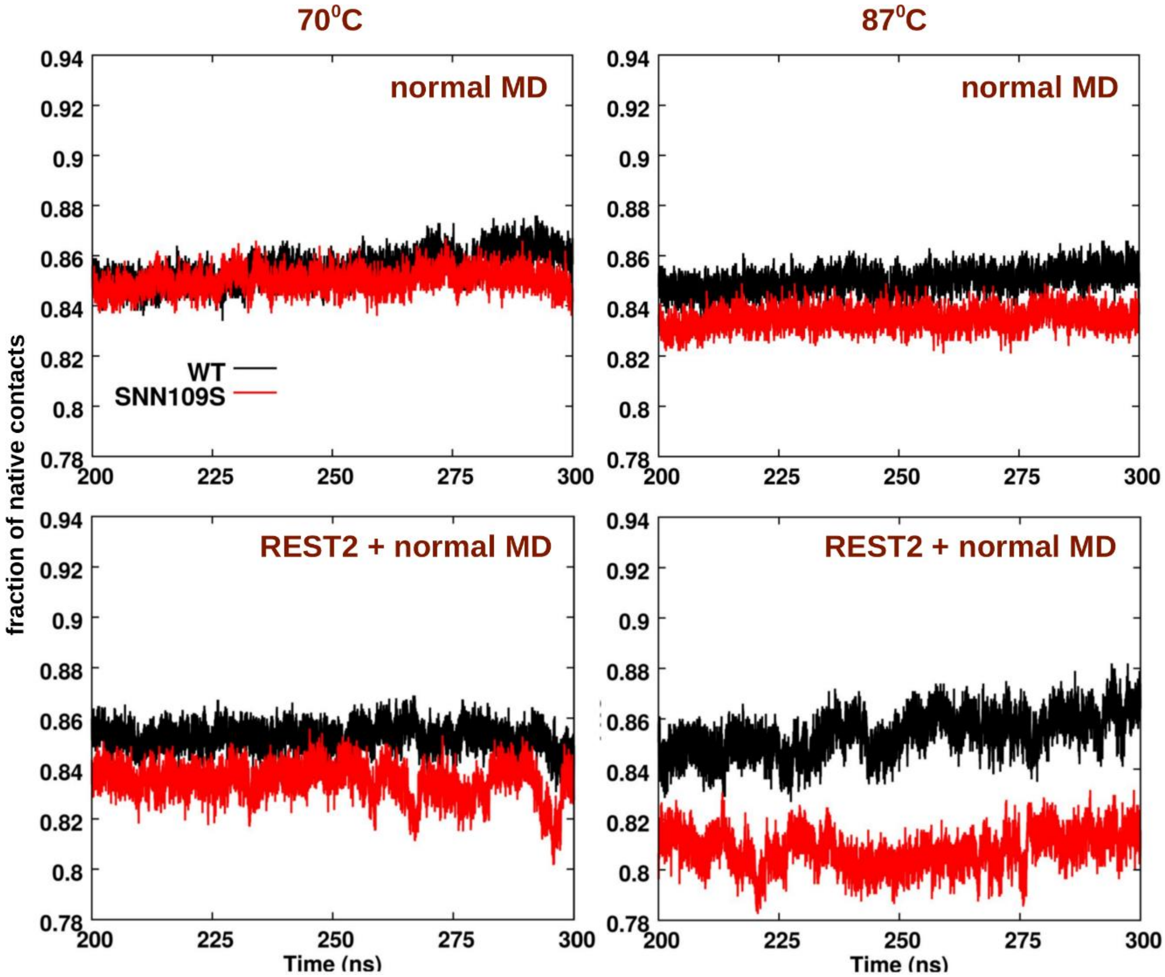


**Supplementary Figure 21:** Comparison of the fraction of native contacts in MjGATase in its WT and SNN109S mutant forms between only normal MD (Run A, see Supplementary Figure 16) and combined REST2 and normal MD approach (Run B).

#### d) Validation of well-tempered metadynamics (WT-MTD) simulations

All the calculations to validate the convergence of WT-MTD simulations and the corresponding free energy profiles were performed (Supplementary Figure 22).

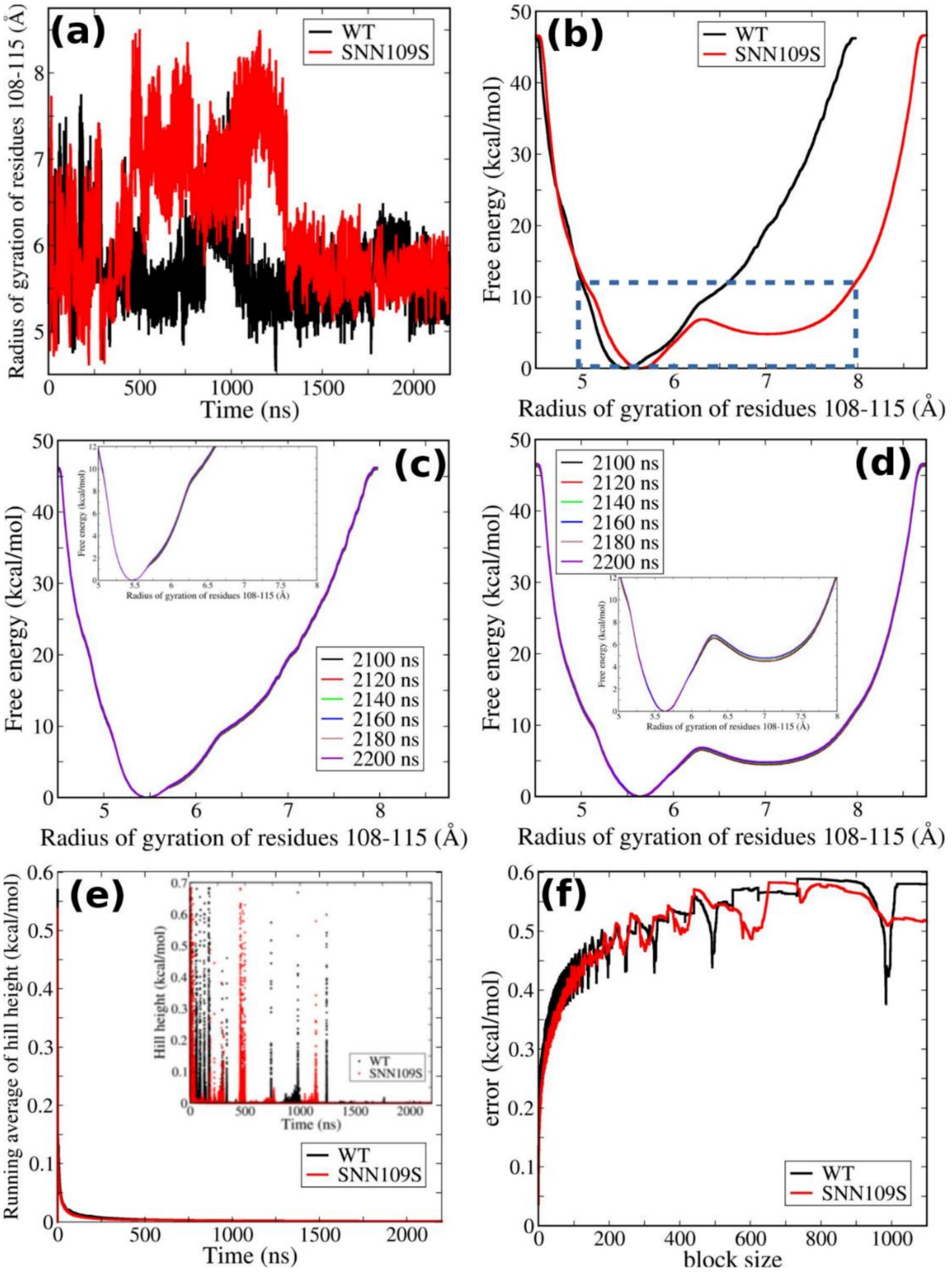


**Supplementary Figure 22:** Validation of well-tempered metadynamics (WT-MTD) simulations. (a) Evolution of the collective variable (CV: radius of gyration of residues near succinimide, i. e., residues 108-115) over simulation time. (b) The full free energy profile generated after 2200 ns of WT-MTD simulation. The important part of this profile is shown within the blue dashed rectangle and the same has been presented in Figure 5a of the main text. (c) Free energy profiles for MjGATase_WT at an interval of 20 ns for the last 100 ns of the simulation (2100-2200 ns) shows that the profile remains almost unchanged over the last 100 ns. The important part is zoomed in the inset. (d) Similar plot like (c), but for MjGATase_SNN109S mutant. (e) Running average of the biasing Gaussian hill height over simulation time. Plot in inset shows the instantaneous hill height. (f) Error estimated from the block averaging(46) shows the convergence of the WT-MTD simulations.

#### ii) Force Field Parameters for Succinimide

#### GROMOS54a7 FF parameters for capped (Ac-SNN-NMe) and uncapped (SNN) succinimide

```
[ Ac-SNN-NMe ]
 [ atoms ]
; name  type    charge      charge group
  CA1   CH3    0.10000    0
    C     C    0.19000    0
    O     O   -0.45000    0
   N3     N   -0.15000    0
    H     H    0.31000    0
   C2     C    0.45000    1
   O2     O   -0.45000    1
   C3   CH1    0.00000    2
   C4  CH2r    0.00000    2
   C5     C    0.45000    3
   O5     O   -0.45000    3
   N1     N    0.00000    4
  CA2   CH3    0.00000    4

 [ bonds ]
;    ai     aj   gromos type
  CA1      C   gb_27
    C      O   gb_5
    C     N3   gb_10
   N3      H   gb_2
   N3     C3   gb_21
   C2     C3   gb_27
   C2     O2   gb_5
   C3     C4   gb_27
   C4     C5   gb_27
   C5     O5   gb_5
   C2     N1   gb_10
   C5     N1   gb_10
   N1    CA2   gb_21

 [ angles ]
;    ai     aj    ak     gromos type
  CA1      C     O     ga_30
  CA1      C    N3     ga_19
    O      C    N3     ga_33
    C     N3     H     ga_32
    C     N3    C3     ga_31
    H     N3    C3     ga_18
   N3     C3    C4     ga_13
   N3     C3    C2     ga_13
   C2     C3    C4     ga_13
   C3     C2    N1     ga_19
   C3     C2    O2     ga_30
   O2     C2    N1     ga_33
```

| | | | |
|---|---|---|---|
| C3 | C4 | C5 | ga_13 |
| C4 | C5 | N1 | ga_19 |
| C4 | C5 | O5 | ga_30 |
| O5 | C5 | N1 | ga_33 |
| C2 | N1 | C5 | ga_21 |
| C2 | N1 | CA2 | ga_31 |
| C5 | N1 | CA2 | ga_31 |

[ improper dihedrals ]

| ; ai | aj | ak | al | gromos type |
|---|---|---|---|---|
| C | CA1 | N3 | O | gi_1 |
| N3 | C | C3 | H | gi_1 |
| C3 | N3 | C2 | C4 | gi_2 |
| C2 | C3 | N1 | O2 | gi_1 |
| C5 | C4 | N1 | O5 | gi_1 |
| N1 | C2 | CA2 | C5 | gi_1 |

[ proper dihedrals ]

| ; ai | aj | ak | al | gromos type |
|---|---|---|---|---|
| C2 | C3 | C4 | C5 | gd_34 |
| N1 | C2 | C3 | C4 | gd_40 |
| C5 | N1 | C2 | C3 | gd_14 |
| C2 | N1 | C5 | C4 | gd_14 |
| N1 | C5 | C4 | C3 | gd_40 |
| CA2 | N1 | C2 | O2 | gd_14 |
| CA2 | N1 | C5 | O5 | gd_14 |
| O2 | C2 | C3 | C4 | gd_40 |
| O5 | C5 | C4 | C3 | gd_40 |
| C2 | C3 | N3 | H | gd_39 |
| C2 | C3 | N3 | C | gd_39 |
| C4 | C3 | N3 | H | gd_39 |
| C4 | C3 | N3 | C | gd_39 |
| C3 | C2 | N1 | CA2 | gd_14 |
| C4 | C5 | N1 | CA2 | gd_14 |
| N3 | C3 | C2 | N1 | gd_40 |
| C2 | N1 | C5 | O5 | gd_14 |
| C5 | N1 | C2 | O2 | gd_14 |
| N3 | C3 | C2 | O2 | gd_40 |
| N3 | C3 | C4 | C5 | gd_34 |
| O | C | N3 | C3 | gd_14 |
| CA1 | C | N3 | C3 | gd_14 |
| O | C | N3 | H | gd_14 |
| CA1 | C | N3 | H | gd_14 |

Note that the nitrogen atom within the succinimide ring is actually not from SNN109 residue, rather it is from the residue next to it (i. e., D110) (Supplementary Figure 10). So, this nitrogen atom is not included within the force field parameters of (uncapped) SNN residue (shown below), rather it is considered within D110.

```
[ SNN ]
 [ atoms ]
; name  type   charge      charge group
    N      N   -0.31000   0
    H      H    0.31000   0
   CA   CH1     0.00000   2
   C4  CH2r     0.00000   2
   C5      C    0.45000   3
   O5      O   -0.45000   3
    C      C    0.45000   1
    O      O   -0.45000   1

 [ bonds ]
;   ai    aj     gromos type
    N      H    gb_2
    N     CA    gb_21
    C     CA    gb_27
    C      O    gb_5
   CA    C4    gb_27
   C4    C5    gb_27
   C5    O5    gb_5
    C    +N    gb_10
   C5    +N    gb_10

 [ angles ]
;   ai     aj    ak    gromos type
   -C      N     H    ga_32
   -C      N    CA    ga_31
    H      N    CA    ga_18
    N     CA    C4    ga_13
    N     CA     C    ga_13
    C     CA    C4    ga_13
   CA      C    +N    ga_19
   CA      C     O    ga_30
    O      C    +N    ga_33
   CA     C4    C5    ga_13
   C4     C5    +N    ga_19
   C4     C5    O5    ga_30
   O5     C5    +N    ga_33
    C     +N    C5    ga_21

 [ impropers dihedrals ]
;   ai    aj    ak    al    gromos type
    N    -C    CA     H    gi_1
   CA     N     C    C4    gi_2
```

```
   C     CA   +N      O     gi_1
  C5     C4   +N    O5     gi_1

 [ proper dihedrals ]
;   ai      aj      ak     al      gromos type
    C     CA     C4    C5     gd_34
    O      C     CA    C4     gd_40
   O5    C5     C4    CA     gd_40
   -C      N    CA     C      gd_44
   -C      N    CA     C      gd_43
    N     CA      C    +N      gd_45
    N     CA      C    +N      gd_42
    N     CA    C4    C5     gd_34
  -CA    -C      N    CA    gd_14
   CA      C    +N    C5     gd_14
```

field. *J Am Chem Soc* 137(7):2695–2703.

30. Abel R, Wang L, Harder ED, Berne BJ, Friesner RA (2017) Advancing Drug Discovery through Enhanced Free Energy Calculations. *Acc Chem Res* 50(7):1625–1632.
31. Bhati AP, Wan S, Coveney P V. (2019) Ensemble-Based Replica Exchange Alchemical Free Energy Methods: The Effect of Protein Mutations on Inhibitor Binding. *J Chem Theory Comput* 15:1265–1277.
32. Cong X, Cremer C, Nachreiner T, Barth S, Carloni P (2016) Engineered human angiogenin mutations in the placental ribonuclease inhibitor complex for anticancer therapy: Insights from enhanced sampling simulations. *Protein Sci*:1451–1460.
33. Bonomi M, et al. (2009) PLUMED: A portable plugin for free-energy calculations with molecular dynamics. *Comput Phys Commun* 180(10):1961–1972.
34. Tribello GA, Bonomi M, Branduardi D, Camilloni C, Bussi G (2014) PLUMED 2: New feathers for an old bird. *Comput Phys Commun* 185(2):604–613.
35. Barducci A, Bussi G, Parrinello M (2008) Well-tempered metadynamics: A smoothly converging and tunable free-energy method. *Phys Rev Lett* 100(2):1–4.
36. Reed AE, Curtiss LA, Weinhold F (1988) Intermolecular Interactions from a Natural Bond Orbital, Donor—Acceptor Viewpoint. *Chem Rev* 88(6):899–926.
37. E. D. Glendening, Reed AE, Carpenter JE, Weinhold F NBO, version 3.1.
38. Frisch, M. J.; Trucks, G. W.; Schlegel, H. B.; Scuseria GE., et al. (2009) Gaussian 09, revision B.01; Gaussian, Inc.: Wallingford, CT, 2009. Available at: https://gaussian.com/.
39. Zhurko G. A. Chemcraft - graphical program for visualization of quantum chemistry computations. https://chemcraftprog.com.
40. Humphrey W, Dalke A, Schulten K (1996) VMD: Visual Molecular Dynamics. *J Mol Graph* 14:33–38.
41. L DeLano W (2002) The PyMOL Molecular Graphics System (2002) DeLano Scientific, Palo Alto, CA, USA. http://www.pymol.org.
42. Daura X, Gademann K, Jaun B, Gunsteren WF Van, Mark AE (1999) Peptide Folding: When Simulation Meets Experiment. *Angew Chem Int Ed*:236–240.
43. Petrov D, Margreitter C, Grandits M, Oostenbrink C, Zagrovic B (2013) A Systematic Framework for Molecular Dynamics Simulations of Protein Post-Translational Modifications. *PLoS Comput Biol* 9(7). doi:10.1371/journal.pcbi.1003154.
44. Margreitter C, Petrov D, Zagrovic B (2013) Vienna-PTM web server: a toolkit for MD simulations of protein post-translational modifications. *Nucleic Acids Res* 41(Web Server issue):422–426.
45. Wang L, Friesner RA, Berne BJ (2011) Replica exchange with solute scaling: A more

efficient version of replica exchange with solute tempering (REST2). *J Phys Chem B* 115(30):9431–9438.

46. Branduardi D, Bussi G, Parrinello M (2012) Metadynamics with adaptive gaussians. *J Chem Theory Comput* 8(7):2247–2254.
