## Supplementary Movies_legend for "Structural basis for the hyperthermostability of an archaeal glutaminase induced by post-translational succinimide formation"

**Supplementary Movie 1: succinimide_stability_WT_27.** Reduced flexibility of the side chain carboxylate of D110 and CO of E108 as visualized from MD simulation

**Supplementary Movie 2:** SNNloop_metaD_WT_27. The movie shows that the succinimide containing loop (residues 108-115) remains folded in MjGATase_WT during the well-tempered metadynamics simulation with the radius of gyration of succinimide loop as the reaction coordinate at 27 °C.

**Supplementary Movie 3:** SNNloop_metaD_SNN109S_27. The movie shows that the loop region from 108-115 in MjGATase_SNN109S can switch between folded and unfolded states during the well-tempered metadynamics simulation with the radius of gyration of residues 108-115 as the reaction coordinate at 27 °C. This indicates that the loop in the mutant protein can unfold at higher temperatures.

**Supplementary Movie 4:** distal_beta-sheet_WT_27. 200 - 300 ns timeframe from REST2 simulation on the extended β-hairpin (residues 86-136) of MjGATase_WT at 27 °C.

**Supplementary Movie 5:** distal_beta-sheet_WT_87. 200 - 300 ns timeframe from REST2 simulation on the extended β-hairpin (residues 86-136) of MjGATase_WT at 87 °C.

**Supplementary Movie 6:** distal_beta-sheet_SNN109S_27. 200 - 300 ns timeframe from REST2 simulation on the extended β-hairpin (residues 86-136) of MjGATase_SNN109S at 27 °C.

**Supplementary Movie 7:** distal_beta-sheet_SNN109S_87. 200 - 300 ns timeframe from REST2 simulation on the extended β-hairpin (residues 86-136) of MjGATase_SNN109S at 87 °C.

**Supplementary Movie 8:** distal_beta-sheet_SNN109P_27. 200 - 300 ns timeframe from REST2 simulation on the extended β-hairpin (residues 86-136) of MjGATase_SNN109P at 27 °C.

**Supplementary Movie 9:** distal_beta-sheet_SNN109P_87. 200 - 300 ns timeframe from REST2 simulation on the extended β-hairpin (residues 86-136) of MjGATase_SNN109P at 87 °C.
